# Defining composition and function of the rhizosphere microbiota of barley genotypes exposed to growth-limiting nitrogen supplies

**DOI:** 10.1101/605204

**Authors:** Rodrigo Alegria Terrazas, Senga Robertson-Albertyn, Aileen Mary Corral, Carmen Escudero-Martinez, Rumana Kapadia, Katharin Balbirnie-Cumming, Jenny Morris, Pete E Hedley, Matthieu Barret, Gloria Torres, Eric Paterson, Elizabeth M Baggs, James Abbott, Davide Bulgarelli

## Abstract

The microbiota populating the rhizosphere, the interface between roots and soil, can modulate plant growth, development and health. These microbial communities are not stochastically assembled from the surrounding soil but their composition and putative function are controlled, at least partially, by the host plant. Here we use the staple cereal barley as a model to gain novel insights into the impact of differential applications of nitrogen, a rate-limiting step for global crop production, on the host genetic control of the rhizosphere microbiota. Using a high-throughput amplicon sequencing survey, we determined that nitrogen availability for plant uptake is a factor promoting the selective enrichment of individual taxa in the rhizosphere of wild and domesticated barley genotypes. Shotgun sequencing and metagenome-assembled genomes revealed that this taxonomic diversification is mirrored by a functional specialisation, manifested by the differential enrichment of multiple GO-terms, of the microbiota of plants exposed to nitrogen conditions limiting barley growth. Finally, a plant soil feedback experiment revealed that the host control on the barley microbiota underpins the assembly of a phylogenetically diverse group of bacteria putatively required to sustain plant performance under nitrogen-limiting supplies. Taken together, our observations indicate that under nitrogen conditions limiting plant growth, plant-microbe and microbe-microbe interactions fine-tune the host genetic selection of the barley microbiota at both taxonomic and functional levels. The disruption of these recruitment cues negatively impacts plant growth.

**Importance:** The microbiota inhabiting the rhizosphere, the thin layer of soil surrounding plant roots, can promote the growth, development, and health of their host plants. Previous research indicated that differences in the genetic composition of the host plant coincide with differences in the composition of the rhizosphere microbiota. This is particularly evident when looking at the microbiota associated to input-demanding modern cultivated varieties and their wild relatives, which have evolved under marginal conditions. However, the functional significance of these differences remains to be fully elucidated. We investigated the rhizosphere microbiota of wild and cultivated genotypes of the global crop barley and determined that nutrient conditions limiting plant growth amplify the host control on microbes at the root-soil interface. This is reflected in a plant- and genotype-dependent functional specialisation of the rhizosphere microbiota which appears required for optimal plant growth. These findings provide novel insights into the significance of the rhizosphere microbiota for plant growth and sustainable agriculture

## Introduction

To sustainably enhance global food security, innovative strategies to increase crop production while preserving natural resources are required (1-3). Capitalising on the microbial communities thriving in association with plants, collectively referred to as the plant microbiota (4, 5), has been identified as one of these innovative strategies (6). For instance, members of the microbiota populating the rhizosphere, the interface between roots and soil, can provide their plant host with access to mineral nutrients and protection against abiotic and biotic stresses (7). Thus, applications of the plant microbiota have the potential to integrate and progressively replace non-renewable inputs in crop production (8).

This potential is of particular interest for alternative to nitrogen (N) applications to staple crops, such as cereals, as approximately 50% of applied fertilisers are lost either in the atmosphere or in groundwater (9, 10). largely as a consequence of microbial denitrification and nitrification processes. Soil microbes can contribute to the release of nitrogen from soil organic matter (SOM) for plant uptake (11). These mineralisation processes are estimated to contribute more than 50% of crop nitrogen (12), even in intensively fertilised systems, and are the fundamental basis of sustained plant productivity in uncultivated soils, as typically more than 90% of soil N is present in organic forms (13). The importance of the plant in influencing these microbial mineralisation processes has been highlighted by the phenomenon of rhizosphere priming effects (14), where root release of organic compounds impacts rates of SOM decomposition and nitrogen mobilisation (15) Therefore, elucidating of the relationships between rhizosphere microbiota composition and nitrogen availability for plant uptake can be a key towards sustainable crop production (16).

The composition of the rhizosphere microbiota is driven, at least in part, by the genetics of its host plants (4, 17). In turn, the processes of domestication and breeding selection, which progressively differentiated wild ancestors from modern, ‘elite’, cultivated varieties (18) modulated plant’s capacity of shaping the microbiota thriving at the root-soil interface (19, 20). As crop wild relatives have evolved in marginal soils, i.e., not exposed to synthetic fertilisers, their microbiota may be equipped with beneficial functions for sustainable agriculture (7, 21). Despite that the impact of plant domestication on the rhizosphere microbiota has been studied in multiple plant species (22-27), the significance of microbial diversification between wild and cultivated plant genotypes of the same species remains to be fully elucidated (21, 28).

Barley (*Hordeum vulgare*), the fourth most cultivated cereal worldwide (29), represents an attractive model to investigate the host genetic control of the rhizosphere microbiota within a framework of plant domestication. For instance, we previously demonstrated that domesticated (*H. vulgare* ssp. *vulgare*) and wild (*H. vulgare* ssp. *spontaneum*) barley genotypes host microbiotas of contrasting composition (30, 31). More recently, we gathered novel insights into the genetic basis of this host-mediated microbiota diversification (32-34). In parallel, investigations targeting specific microbial genes indicated that barley plants may exert a control on microbes underpinning the nitrogen biogeochemical cycle (35) and that this effect is dependent on community composition (36). However, it is unclear how genetic differences between wild and domesticated genotypes may impact on the composition and function of the rhizosphere microbiota of plants exposed to contrasting nitrogen supplies, in particular the ones limiting plant growth.

To address this knowledge gap, in this investigation we used barley as an experimental model and state-of-the art sequencing approaches to test three interconnected hypotheses. First, we hypothesised that the host control on rhizosphere bacteria is modulated by, and responds to, nitrogen availability for plant uptake. Specifically, we anticipated that differences in microbiota composition among barley genotypes is maximal under limiting nitrogen supplies, when plants rely on their microbiota for N-cycling processes to support optimal growth. We further hypothesised that, under conditions limiting barley growth, plant’s reliance on the rhizosphere microbes will be manifested by a functional diversification mediated, at least partially, by the host genotype. Finally, we hypothesised that that these distinct structural and functional configurations of the microbiota contributed to differential plant growth responses.

## Results

### Nitrogen conditions limiting plant growth amplify the host effect on the barley rhizosphere microbiota

To gain insights into the role played by nitrogen availability for plant uptake on the composition of the barley bacterial microbiota, we selected one reference barley cultivar, Morex (hereafter ‘Elite’), and two wild genotypes from the B1K collection (37), B1K-12 and B1K-31 (hereafter ‘Desert’ and ‘North’ respectively). The rationale for this choice was two-pronged. First, we previously characterised these genotypes for their capacity for recruiting distinct microbiotas and genetic relatedness (30, 31). Second, the wild genotypes are representative of the two main barley ecotypes identified in the Southern Levant as drivers of plant’s adaptation to the environment (38, 39). Consequently, and despite the limited number, these genotypes may capture the “extremes” of the evolutionary pressure on the host recruitment cues of the barley microbiota. Plants were grown under glasshouse conditions (Methods) in an agricultural soil previously used for microbiota investigations and designated ‘Quarryfield’ (31, 33, 34, 40). Pots containing the individual genotypes, and unplanted soil controls (hereafter ‘Bulk’), were supplemented with three modified Hoagland’s solution preparations (41) containing all essential macro and micronutrients and three levels of mineral nitrogen (Table S1): the optimum required for barley growth (N100%), a quarter of dosage (N25%) or no nitrogen (N0%). At early stem elongation (Figure S1), which represents the onset of maximum nitrogen uptake for small grain cereals (42) plants were harvested, and total DNA preparations were obtained from rhizosphere and unplanted soil specimens. In parallel, we determined aboveground plant biomass, plant nitrogen content in leaves as well as concentrations of ammonium (NH_4_ ^+^) and nitrate (NO_3_ ^-^) in rhizosphere and unplanted soil samples.

We observed that plant performance was affected by the N application: aboveground biomass and plant nitrogen content were significantly lower at N0% compared to N100%, with N25% yielding intermediate values (Kruskal-Wallis test followed by Dunn post-hoc test, individual *P* values <0.05, FDR corrected, Figure 1) compatible with a nitrogen-deficiency status for barley growth. Likewise, the residual nitrogen in the rhizosphere at the completion of the experiments, measured as a concentration of ammonium and nitrate respectively, displayed a significant decrease in the values recorded for N100% to N25% and from this latter to N0%. (Kruskal-Wallis test followed by Dunn post-hoc test, individual *P* values <0.05, FDR corrected, Figure 1).

**Figure 1:**
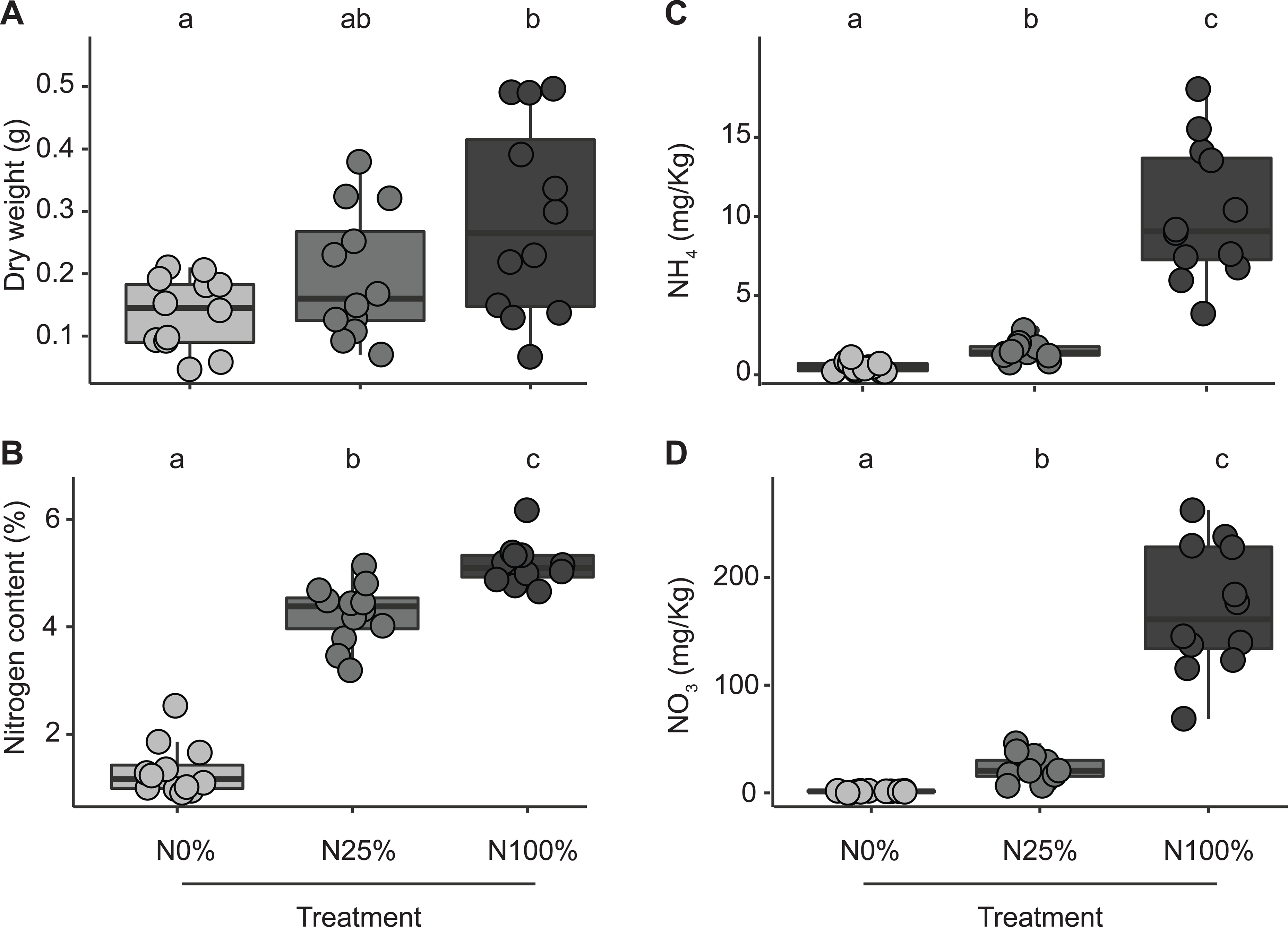
The nitrogen content of ‘Quarryfield’ soil limits barley growth. Cumulative data gathered at early stem elongation in the tested barley genotypes subjected to three nitrogen fertilization treatments (N0%, N25% and N100%) as indicated in the x-axis of each panel. Individual dots depict individual biolgical replicates. A) aboveground biomass of the tested plants. B) nitrogen content in the aboveground tissues of the tested plants. Residual concentration of C) ammonium and D) nitrate retrieved from rhizospheric soil at the time of sampling. Lowercase letters denote significant differences at *P* value <0.05 in a Kruskal–Wallis non-parametric analysis of variance followed by Dunn’s post hoc test.

In parallel, we generated a 16S rRNA gene amplicon sequencing library from the obtained rhizosphere and unplanted soil controls and we identified 26,411 individual Amplicon Sequencing Variants (ASVs) accruing from 6,097,808 sequencing reads. After pruning *in silico* ASVs representing either host (i.e., plastid- or mitochondrial-derived sequences) or environmental contaminations, as well as low count ASVs, 5,035,790 reads were retained, representing over the 82% of the initial dataset. Canonical Analysis of Principal Coordinates (CAP) differentiated bulk soil from rhizosphere profiles as evidenced by a segregation of either class of samples along the axis accounting for the largest source of variation (Figure 2A).

**Figure 2.**
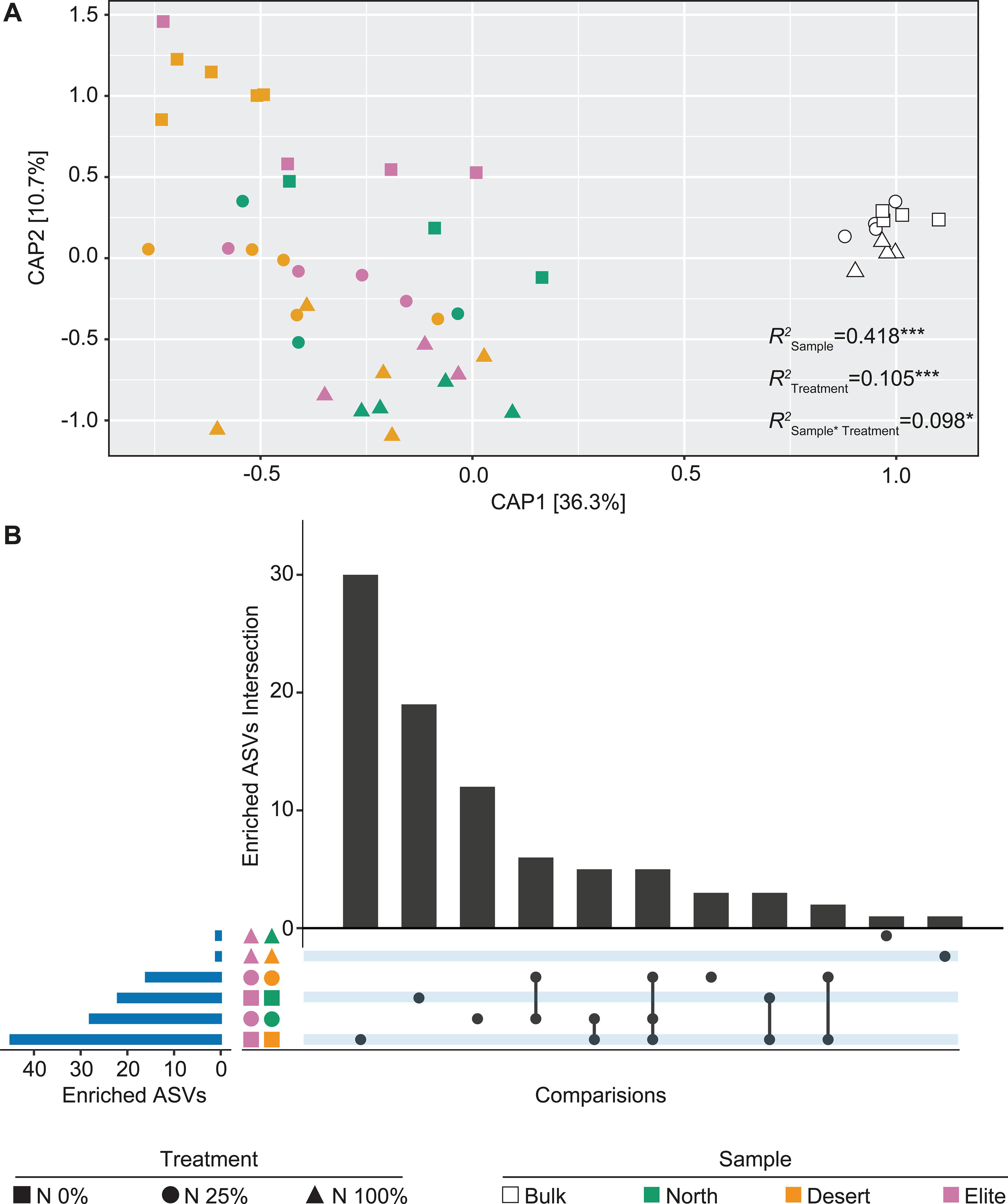
Nitrogen availability modulates the host genetic control of the rhizosphere bacterial microbiota. A) Canonical analysis of Principal Coordinates computed on Bray-Curtis dissimilarity matrix. Individual shapes in the plot denote individual biological replicates whose colour and shape depict sample type and nitrogen treatment, respectively, as indicated in the bottom part of the figure. Numbers in the plots depict the proportion of variance (R2) explained by the factors ‘Sample’, ‘Treatment’ and their interactions, respectively. Asterisks associated to the R2 value denote its significance, *P* value ‘Sample’ = 0.0002, *P* value ‘Treatment’ = 0.0004, *P* value ‘Sample * Treatment’ = 0.0434; Adonis test 5,000 permutations. B) Horizontal blue bars denote the number of ASVs differentially enriched (Wald test, Individual *P* values <0.05, FDR corrected between the elite and two wild barley genotypes at different nitrogen treatments as recapitulated by the shape and colour scheme. Vertical bars depict the number of differentially enriched ASVs unique for or shared among two or more pair-wise comparisons highlighted by the interconnected dots underneath the vertical bars.

Furthermore, we observed a “gradient” along the axis accounting for the second source of variation aligned with the treatment effect, in particular for rhizosphere samples (Figure 2A). The sample effect, i.e., either bulk soils or the rhizosphere of the individual genotypes, exerted the primary impact on the bacterial communities thriving at the root-soil interface (Permanova, *R*^2^= 0.418, *P* value = 0.0002, 5,000 permutations, Figure 2A) followed by the nitrogen treatment effect (Permanova, *R*^2^= 0.105, *P* value =0.0004; 5,000 permutations, Figure 2B) and their interaction term (Permanova, *R*^2^= 0.098, *P* value =0.0380; 5,000 permutations, Figure 2B).

To further examine the impact of the treatment on the abundance of individual prokaryotic ASVs underpinning host-mediated diversification, we performed a set of pair-wise comparisons between barley genotypes at the different N levels. We observed that N0% was associated with the largest number of differentially recruited ASVs, while higher N levels progressively obliterated recruitment differences among genotypes (Wald test, Individual *P* values < 0.05, FDR corrected, Figure 2B). Of note, and congruent with previous experiments conducted in the same soil (31), the pair ‘Elite’-‘Desert’ yielded the highest number of differentially recruited ASVs (Figure 2B).

Taken together, these observations indicate that nitrogen availability for plant uptake is a factor in *a*) modulating the microhabitat- and genotype-dependent recruitment cues of the barley bacterial microbiota, by *b*) promoting the selective enrichment of individual taxa in the rhizosphere and, *c*) whose magnitude is maximised when no nitrogen is applied to the system.

The metabolic potential of the rhizosphere microbiota exposed to nitrogen conditions limiting barley growth

We generated over 412 million paired-end metagenomic reads from 12 additional samples to gain insights into the functional significance of microbiota diversification in plants exposed to nitrogen conditions limiting barley growth. These represented three biological replicates each of Bulk soil and the rhizosphere of ‘Elite’, ‘North’ and ‘Desert’ exposed to the N0% treatment. Upon in-silico removal of low-quality sequences and sequences matching the barley genome, likely representing a “host contamination” (Figure S2), taxonomic classification of the sequencing reads at kingdom level revealed that Bacteria outnumbered Fungi by two orders of magnitude, regardless of the sample investigated (Figure 3A). Closer inspection of the data classified within the kingdom fungi revealed no significant differences among samples for sequences assigned to the class Glomeromycetes, which we used as a proxy for the extra-radical mycelium of Arbuscular Mycorrhizal Fungi (AMF; Wald test, Individual *P* values > 0.05, FDR corrected, Figure 3B).

**Figure 3.**
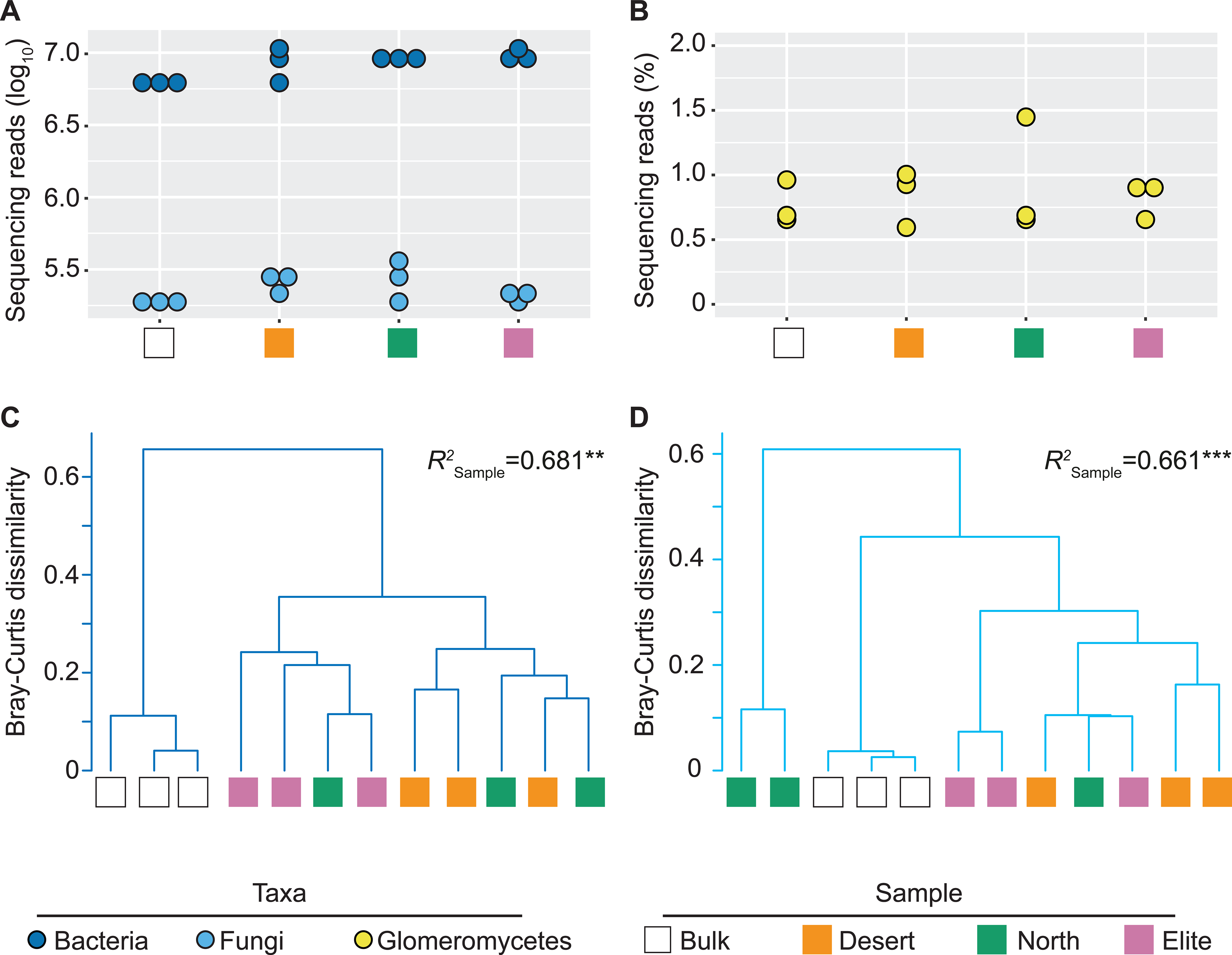
Bacteria dominate the metagenome of barley plants exposed to limiting nitrogen supplies. Dots depict sequencing reads assigned to A) Bacteria and Fungi or B) proportion of fungal sequencing reads classified as ‘Glomeromycetes’ in the individual replicated of the metagenomic survey in the indicated samples. In C) and D) cluster dendrograms constructed using Bray-Curtis dissimilarity matrices of the metagenomic sequencing reads (counts per million) assigned to family level in Bacteria and Fungi, respectively. Individual shapes denote individual biological replicates whose colour depict sample type as indicated in the bottom part of the figure. Numbers associated to each dendrogram depict the proportion of variance (R2) explained by the factor ‘Sample’, in Bacteria or Fungi, respectively. Asterisks associated to the R2 value denote it significance, *P* value ‘Sample’ in Bacteria = 0.0012; *P* value ‘Sample’ in Fungi = 0.0004; Adonis test 5,000 permutations.

Although the separation between replicates of the same genotype, in particular the ‘Elite’-‘Desert’ pair, was manifested exclusively when looking at bacteria, we identified a comparable effect of the sample type on composition of both bacterial and fungal communities. For instance, the *R*^2^ computed for normalised relative abundances returned values between 0.66 and 0.68 for the bacterial and fungal component, respectively (Permanova, 5,000 permutations, Individual *P* values < 0.01; Figures 3C and 3D).

Next, we mined the metagenomic dataset for sequencing reads associated with known genes underpinning the nitrogen biogeochemical cycle. We were able to identify genes implicated in processes as diverse as nitrification, denitrification, nitrate reduction as well as synthesis and degradation of nitrogen-containing organic compounds, although the abundances of genes associate to the individual process did not discriminate between barley genotypes (Wald test, Individual *P* values > 0.05, FDR corrected, Figure S3). This suggests that, in the tested conditions, the host control of the nitrogen biogeochemical cycle does not represent the main driver of the functional diversification of the barley rhizosphere microbiota.

This motivated us to further discern the metabolic capacity of barley associated communities, by assembling metagenomic reads and predicting their encoded proteins. Predicted proteins were clustered, resulting in 10,554,104 representative sequences. The representative protein sequences were subjected to functional enrichment analysis to identify GO categories differentially enriched in the barley rhizosphere. We observed a consistent ‘rhizosphere effect’ in the functional potential of the barley microbiota manifested by a spatial separation of plant-associated communities from Bulk soil in an ordination (Figure 4A) sustained by a differential enrichment of multiple GO categories (Figure 4B). Closer inspection of these categories revealed a significant enrichment of multiple GO terms in each of the tested genotypes and the Bulk soil alike (Wald test, Individual *P* values <0.05, FDR corrected, Table 2). In particular, the microbiota associated to ‘Desert’, ‘North’ and the ‘Elite’ genotypes enriched for of GO-terms implicated in carbohydrate metabolic processing; cell adhesion, pathogenesis, response to abiotic stimulus, responses to chemical, protein-containing complex assembly as well as bacterial-type flagellum-dependent cell motility. These enrichments appear congruent with the adaptation of polymicrobial communities to a host capable of providing substrates for microbial growth. Conversely, bulk soil specimens were enriched predominantly for functions implicated photosynthesis and sporulation which are congruent with microbial adaptation to a lack of organic resources such as the case in unplanted soils.

**Figure 4.**
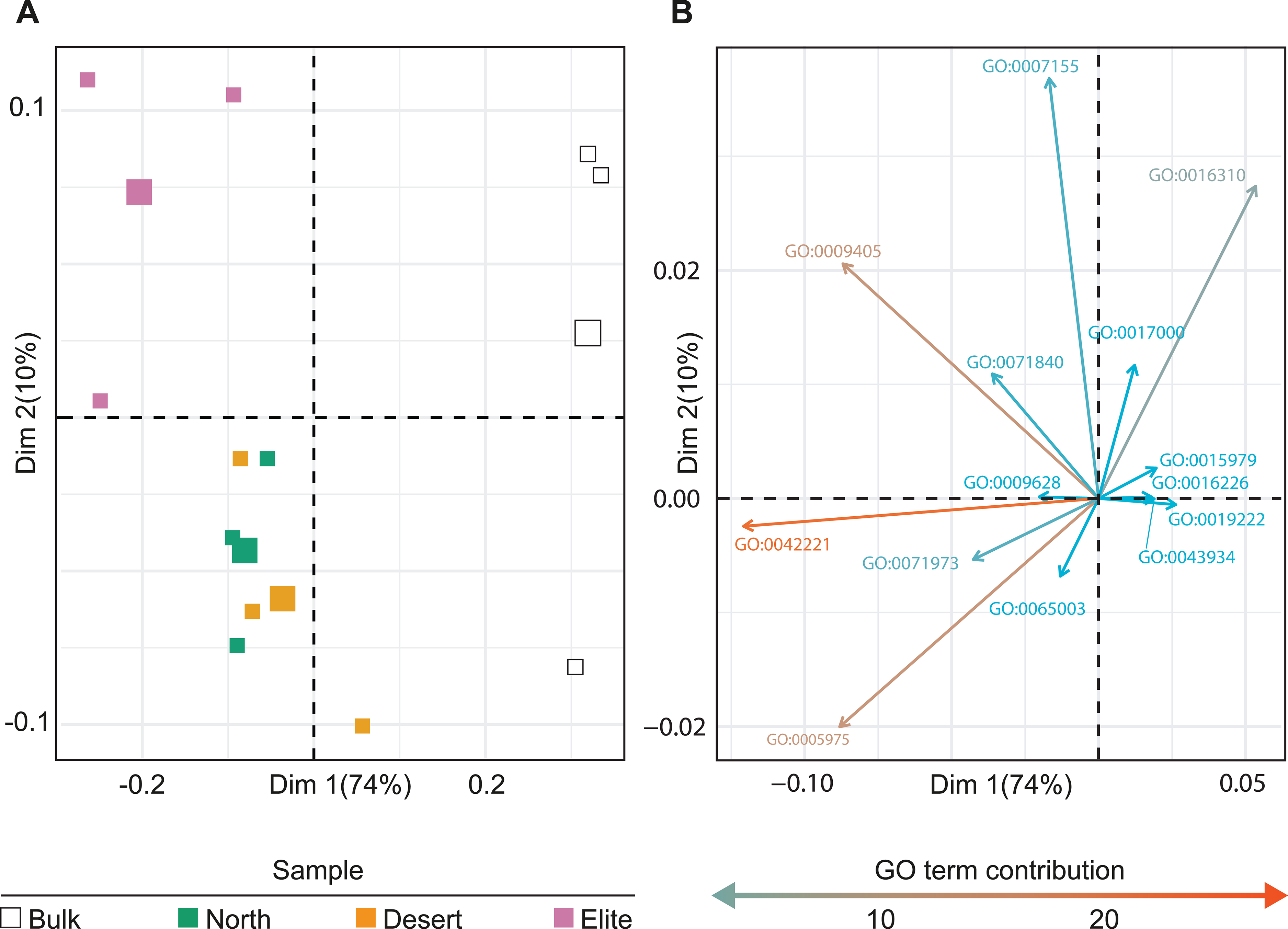
The microhabitat and the host genotype fine-tune the functional potential of the barley microbiota. A) Principal Component Analysis computed on annotated reads mapped to the terms of Gene Ontology Slim database. Individual shapes in the plot denote individual biological replicates whose colour depict sample type as indicated in the bottom of the panel. The largest shape of each sample type indicates the centroid. B) PCA loadings representing the GO Slim terms sustaining the ordination. The top 20 GO Slim terms were filtered for those with a log2 fold-change of > ± 0.2 in at least one comparison (Wald test, Individual *P* values <0.05, FDR corrected). Arrows point at the direction of influence of a given term in the various samples, their length and colour being proportional to the weight they contribute to each PC as indicated in the key underneath the plot.

**Table 1:** Metagenomic Assembly statistics.

|  |  |
| --- | --- |
| Assembly Length | 12841009562 bp |
| Number Contigs | 21005959 |
| Longest Contig | 579848 bp |
| N50 | 627 bp |
| L50 | 5320588 bp |
| Predicted Protein Sequences | 26740734 |
| Protein Sequence Clusters | 10554104 |

**Table 2:** GO Slim Terms with Significantly Differential Abundance.

| Accession | GO Term | Bulk-Desert |  | Bulk-North |  | Bulk-Elite |  | Desert-Elite |  | North-Elite |  | Desert-North |  |
| --- | --- | --- | --- | --- | --- | --- | --- | --- | --- | --- | --- | --- | --- |
|  |  | log2 FC | FDR | log2 FC | FDR | log2 FC | FDR | log2 FC | FDR | log2 FC | FDR | log2 FC | FDR |
| GO:0005975 | carbohydrate metabolic process | 0.23 | $3.59 \times 10^{-07}$ | 0.29 | $2.42 \times 10^{-11}$ | 0.34 | $1.03 \times 10^{-14}$ | $1.05 \times 10^{-01}$ | $3.99 \times 10^{-02}$ | - | NS | $6.37 \times 10^{-02}$ | $8.28 \times 10^{-01}$ |
| GO:0007155 | cell adhesion | -0.06 | $6.62 \times 10^{-01}$ | 0.03 | $8.11 \times 10^{-01}$ | 0.26 | $2.17 \times 10^{-02}$ | $3.23 \times 10^{-01}$ | $1.84 \times 10^{-02}$ | - | NS | $8.97 \times 10^{-02}$ | $9.10 \times 10^{-01}$ |
| GO:0009405 | Pathogenesis | 1.21 | $4.64 \times 10^{-05}$ | 1.25 | $2.65 \times 10^{-05}$ | 1.80 | $2.66 \times 10^{-10}$ | $5.88 \times 10^{-01}$ | $9.05 \times 10^{-02}$ | - | NS | $3.26 \times 10^{-02}$ | $9.79 \times 10^{-01}$ |
| GO:0009628 | response to abiotic stimulus | 1.33 | $4.65 \times 10^{-04}$ | 1.45 | $1.03 \times 10^{-04}$ | 1.70 | $3.20 \times 10^{-06}$ | $3.66 \times 10^{-01}$ | $4.26 \times 10^{-01}$ | - | NS | $1.22 \times 10^{-01}$ | $9.79 \times 10^{-01}$ |
| GO:0015979 | Photosynthesis | -0.84 | $1.18 \times 10^{-04}$ | -0.89 | $4.13 \times 10^{-05}$ | -1.20 | $1.03 \times 10^{-08}$ | $-3.60 \times 10^{-01}$ | $1.69 \times 10^{-01}$ | - | NS | $-4.63 \times 10^{-02}$ | $9.79 \times 10^{-01}$ |
| GO:0016226 | iron-sulfur cluster assembly | -0.16 | $7.53 \times 10^{-06}$ | -0.16 | $1.55 \times 10^{-06}$ | -0.21 | $1.70 \times 10^{-10}$ | $-5.77 \times 10^{-02}$ | $1.69 \times 10^{-01}$ | - | NS | $-8.71 \times 10^{-03}$ | $9.79 \times 10^{-01}$ |
| GO:0016310 | Phosphorylation | -0.14 | $3.24 \times 10^{-02}$ | -0.18 | $6.04 \times 10^{-03}$ | -0.18 | $5.10 \times 10^{-03}$ | $-3.45 \times 10^{-02}$ | $6.95 \times 10^{-01}$ | - | NS | $-3.66 \times 10^{-02}$ | $9.79 \times 10^{-01}$ |
| GO:0017000 | antibiotic biosynthetic process | -0.24 | $2.72 \times 10^{-03}$ | -0.15 | $6.67 \times 10^{-02}$ | -0.15 | $4.81 \times 10^{-02}$ | $8.49 \times 10^{-02}$ | $4.08 \times 10^{-01}$ | - | NS | $9.06 \times 10^{-02}$ | $8.28 \times 10^{-01}$ |
| GO:0019222 | regulation of metabolic process | -0.21 | $2.72 \times 10^{-03}$ | -0.21 | $3.20 \times 10^{-03}$ | -0.36 | $1.13 \times 10^{-07}$ | $-1.42 \times 10^{-01}$ | $8.09 \times 10^{-02}$ | - | NS | $5.68 \times 10^{-03}$ | $9.79 \times 10^{-01}$ |
| GO:0042221 | response to chemical | 0.61 | $7.40 \times 10^{-11}$ | 0.61 | $2.42 \times 10^{-11}$ | 0.76 | $5.61 \times 10^{-17}$ | $1.51 \times 10^{-01}$ | $1.69 \times 10^{-01}$ | - | NS | $5.19 \times 10^{-03}$ | $9.79 \times 10^{-01}$ |
| GO:0043934 | Sporulation | -0.48 | $2.08 \times 10^{-05}$ | -0.57 | $2.62 \times 10^{-07}$ | -0.81 | $4.76 \times 10^{-14}$ | $-3.31 \times 10^{-01}$ | $1.84 \times 10^{-02}$ | - | NS | $-9.21 \times 10^{-02}$ | $9.10 \times 10^{-01}$ |
| GO:0065003 | protein-containing complex assembly | 0.59 | $4.67 \times 10^{-07}$ | 0.60 | $2.62 \times 10^{-07}$ | 0.53 | $3.29 \times 10^{-06}$ | $-5.78 \times 10^{-02}$ | $6.95 \times 10^{-01}$ | - | NS | $7.51 \times 10^{-03}$ | $9.79 \times 10^{-01}$ |
| GO:0071840 | cellular component organization or biogenesis | 0.12 | $1.90 \times 10^{-02}$ | 0.18 | $3.06 \times 10^{-04}$ | 0.25 | $5.59 \times 10^{-07}$ | $1.22 \times 10^{-01}$ | $3.99 \times 10^{-02}$ | - | NS | $5.89 \times 10^{-02}$ | $8.28 \times 10^{-01}$ |
| GO:0071973 | bacterial-type flagellum-dependent cell motility | 0.75 | $3.82 \times 10^{-13}$ | 0.71 | $3.51 \times 10^{-12}$ | 0.86 | $1.70 \times 10^{-17}$ | $1.07 \times 10^{-01}$ | $4.26 \times 10^{-01}$ | - | NS | $-3.68 \times 10^{-02}$ | $9.79 \times 10^{-01}$ |

Significantly differentially abundant GO Slim terms (Benjamini-Hochberg corrected FDR<0.05, Wald test) with log2 fold-change > ±0.20 in at least one comparison. NS=Not Significant.

To gain a finer view of the functional diversification of the bulk and rhizosphere microbiotas, we performed a cluster analysis of individual GO-terms on the top 10 clusters differentiating between samples (Figure S4). For each cluster we determined the significance of individual terms in pair-wise comparisons between bulk soil and rhizosphere samples and, within the latter, between genotypes (Wald test, Individual *P* values <0.05, FDR corrected, Dataset S1). This allowed us to implicate nitrate transporters with functions putatively underpinning multitrophic interactions, such as response to reactive oxygen species and the type VI secretion system. These two functions were also significantly enriched in and differentiating between ‘Elite’ and ‘Desert’ communities (Wald test, Individual *P* values <0.05, FDR corrected, Dataset S1, cluster 6). Conversely, ammonium transporters were identified as a depleted function function in rhizosphere communities (Wald test, Individual *P* values <0.05, FDR corrected, Dataset S1, clusters 5 and 8) and so were functions implicated in phosphate metabolism, including ‘cellular phosphate homeostasis’, ‘negative regulation of phosphate metabolic process’ and ‘phosphate ion transmembrane transport’ (Wald test, Individual *P* values <0.05, FDR corrected, Dataset S1, clusters 5 and 8). The overarching picture emerging in this investigation was that, at the metagenomic resolution we obtained, the major effect on the functional potential of the microbiota is exerted by the microhabitat (i.e., bulk vs. rhizosphere). Conversely, the effect of the host genotype appears confined to a limited number of individual GO-terms and, congruently with the 16S rRNA gene survey, manifested predominantly in the comparison between ‘Elite’ and ‘Desert’ genotypes.

### Genome reconstruction of the bacteria populating the barley root-soil interface

As a first step towards linking structural and functional diversification of the barley microbiota, we attempted to reconstruct genomes of individual bacteria proliferating at the root-soil interface. We assembled the generated metagenomic reads and combined contigs with similar nucleotide composition and differential abundance across samples. This resulted in the reconstruction of 60 Metagenome-Assembled Genomes (MAGs) with a completion of > 50 % according to the presence of a minimal set of essential genes and a proportion of contamination < 10% (Methods). These MAGs were taxonomically affiliated to 12 different bacterial classes and their genome systematically mined for the top 10 GO-terms significantly enriched in the rhizosphere samples compared to Bulk soil controls (Wald test, individual *P* values < 0.05, FDR corrected; Figure 5). Next, we determined co-occurrence patterns between these terms and identified two clusters. One of those linking genomes coding for ‘photosynthesis’, ‘carbohydrate metabolic process’ and ‘iron-sulphur cluster assembly’, while another one linking ‘cellular component organization or biogenesis’, ‘response to chemical’, ‘bacterial-type flagellum-dependent cell motility’ and ‘protein-containing complex assembly’ (Pearson’s correlation, individual *P* value < 0.05; Figure 6). When we interpolated the results of these two analyses, we observed that this second cluster is predominantly represented by MAGs classified as Proteobacteria while the ‘carbohydrate metabolic process’ defining the first cluster was preferentially associated to MAGs classified as Bacteroidetes. For 11 of the 15 MAGs assigned to this phylum the presence of ‘carbohydrate metabolic process’ predicted a significant enrichment of given MAGs in the microbiota of the Elite variety (Figure S5, Wald test, individual *P* value <0.05, FDR corrected). Conversely, among Proteobacterial MAGs, the selected GO terms failed to predict enrichment patterns in a given plant genotype (Figure S5, Wald test, individual *P* value <0.05, FDR corrected).

**Figure 5.**
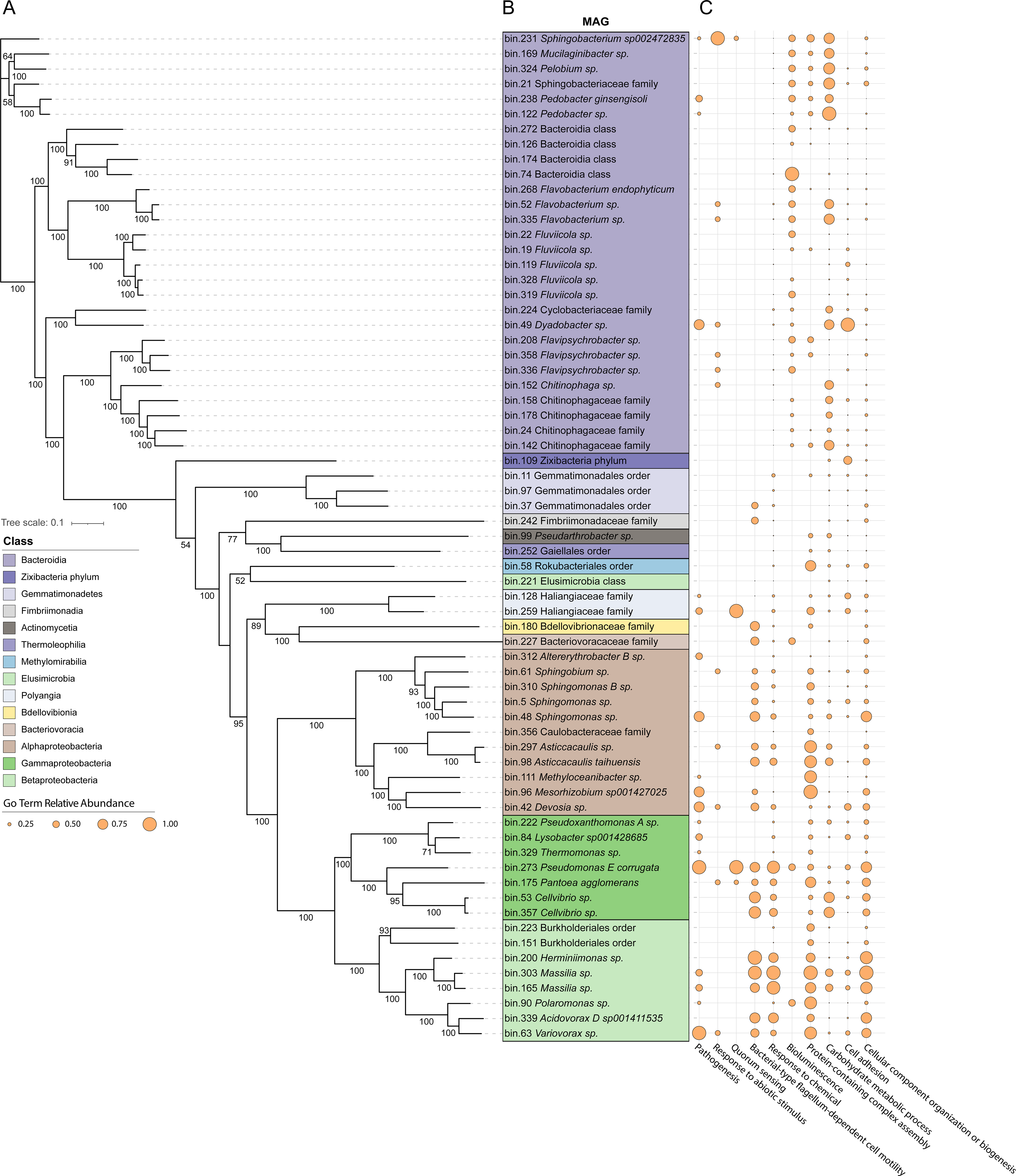
Partitioning of the functional potential of the rhizosphere microbiota among its individual members A) Core gene based phylogenetic tree of the 60 MAGs identified in this study. Branch labels represent bootstrap values (100 bootstrap iterations) B) Taxonomic affiliation of the individual MAGs obtained using GTDB-TK, highlighting colours denote class affiliation as indicated in panel in the left end-side of the figure. C) Distribution of sequences mapping to the top 10 GO Slim categories significantly enriched in the rhizosphere samples compared to Bulk soil controls (Wald test, Individual *P* values <0.05, FDR corrected). The size of the dots denotes the relative abundance of each annotated term in a given genome.

**Figure 6.**
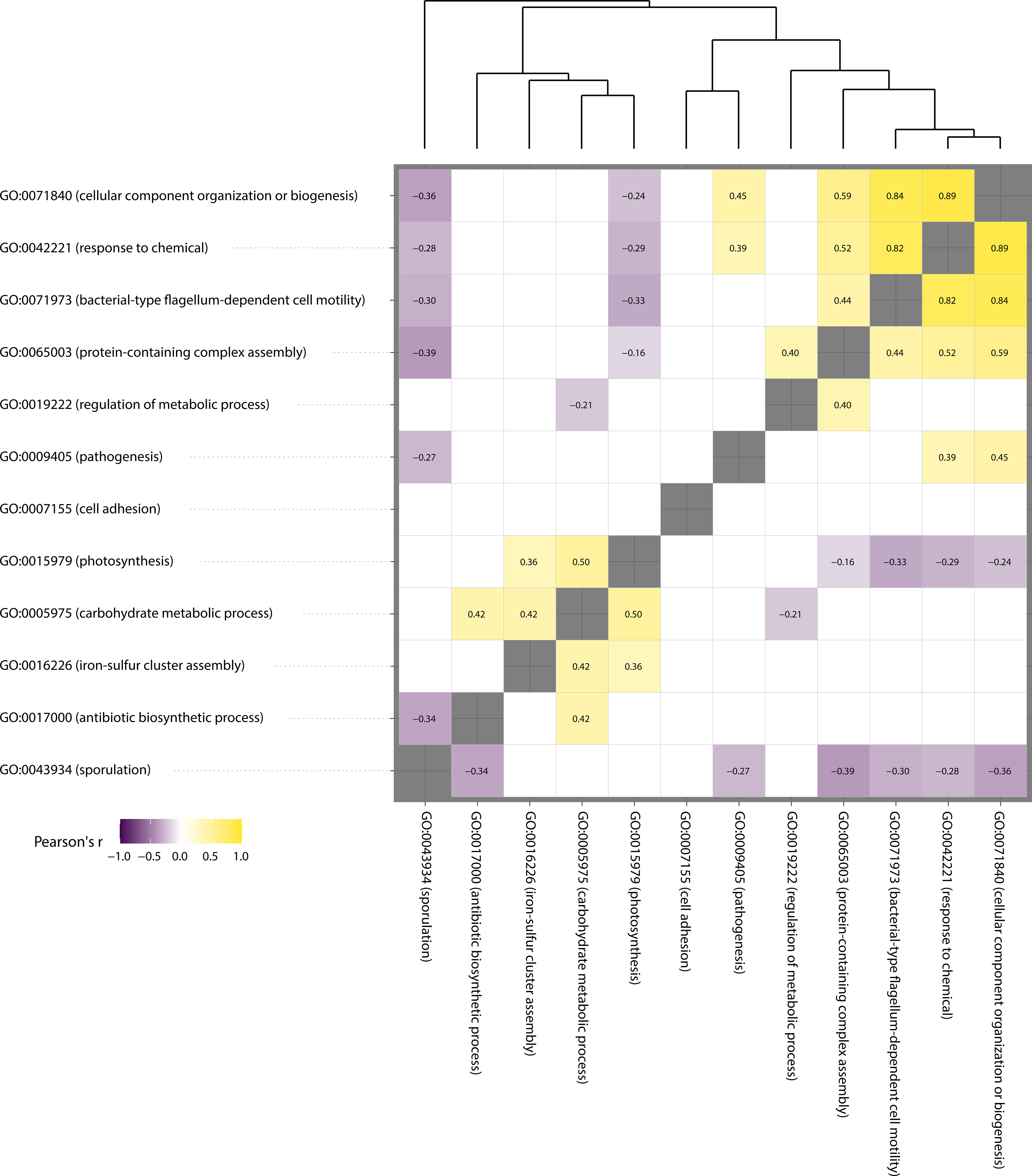
Co-occurrence of individual GO Terms in the barley rhizosphere metagenome. Pair-wise correlation among the abundances of individual GO terms identified in the MAGs. Individual numbers in the plot depict Pearson’s r correlation coefficient. This coefficient is reported for only pair-wise correlations displaying individual *P* values < 0.05.

### A distinct bacterial consortium is associated to optimum barley growth under nitrogen limiting supply

To establish a causal relationship between structural and functional configurations of the rhizosphere microbiota and plant growth, we performed a plant-soil feedback experiment by growing the ‘Elite’ variety in soils previously used for the growth of either a domesticated or wild genotypes amended with a N0% solution (hereafter ‘conditioned soil’). For this analysis we focused on the pair ‘Elite’ – ‘Desert’ as these genotypes displayed the most contrasting microbiota (e.g., Figure 2 and 3). The obtained conditioned soils were used either in their ‘native’ form or subjected to a heat treatment which we hypothesized led to a disruption of the taxonomic and functional configurations of the microbiota (Figure 7A).

**Figure 7.**
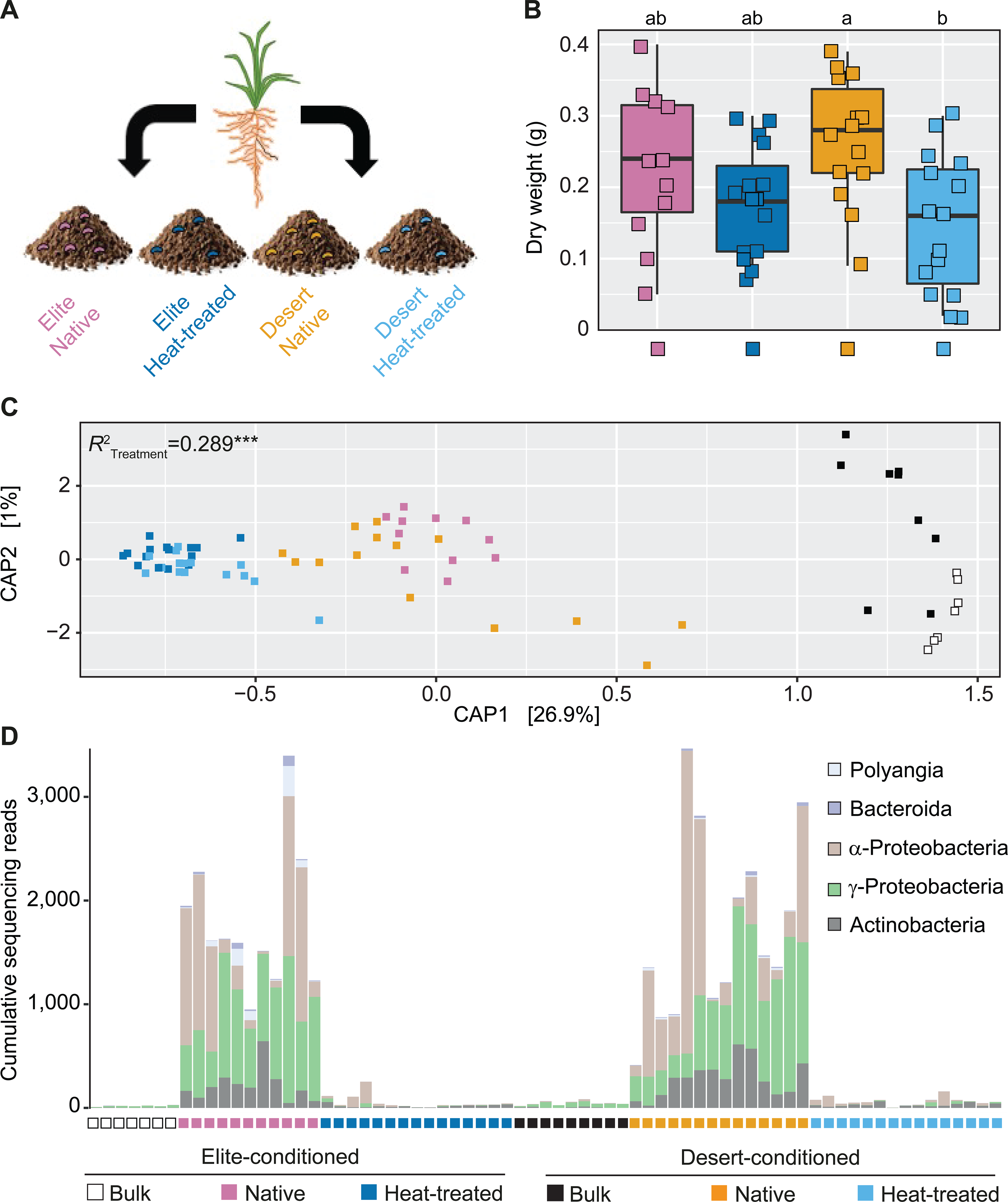
A phylogenetically diverse bacterial consortium is associated to optimum barley growth in plants exposed to nitrogen-limiting conditions. A) Schematic representation of the implemented plant-soil feedback experiments. B) Above-ground biomass of ‘Elite’ barley plants sampled at early stem elongation in conditioned soil in either native or heat-treated form as indicated by colour-coding at the bottom of the figure. Dots depict individual biological replicates and letters denote significant differences at *P* value <0.05 in a two-way analysis of variance followed by a Tukey’s post hoc test. C) Canonical analysis of Principal Coordinates computed on Bray-Curtis dissimilarity matrix. Individual shapes in the plot denote individual biological replicates whose colour depict sample type and treatment. i.e., native or heat-treated, as indicated in the bottom part of the figure. The number in the plot depict the proportion of variance (R2) explained by the factor, ‘Treatment’ while the asterisks define its significance, P value ‘Treatment’ = 0.0004; Adonis test 5,000 permutations. D) Cumulative abundances, expressed as number of sequencing reads, assigned to each of the bacteria significantly enriched in and discriminating between rhizosphere of plants grown in native, conditioned soil versus both Bulk and heat-treated conditioned soils (Wald test, Individual *P* values <0.05, FDR corrected). Each vertical bar corresponds to an individual biological replicate of a sample and treatment depicted underneath the graph. Each segment in the vertical bar depicts the sequencing reads assigned to an individual bacterial ASV, color-coded according to its affiliation at class level.

Plants grown in the ‘heat-treated’ soil displayed a growth deficit when compared to their native counterpart, although these differences were significant only for the Desert-conditioned soil (two-way ANOVA followed by TukeyHSD test, Individual *P* values <0.05; Figure 7B). Closer inspection of 19 chemical and physical parameters characterising the conditioned soils failed to single out a ‘Desert-specific’ parameter. Rather, a limited number of properties explained most of the variance among samples and differentiated between native soil and their heat-treated counterparts, irrespective of the initial genotype used (Statistical values for the individual properties: *P* value < 0.01, *R*^2^ > 0.8; 5,000 permutations; Figure S6). Conversely, qPCR analyses of 16S rRNA gene and ITS copy numbers performed at the end of the cultivation revealed a Desert-mediated impact on the bacterial but not on the fungal communities populating the conditioned soils (Kruskal-Wallis test followed by Dunn post-hoc test, *P* value = 0.039, FDR corrected, Figure S7).

This observation motivated us to gain insights into the taxonomic composition of the bacterial communities inhabiting the conditioned soil. A 16S rRNA gene amplicon library constructed from the samples subjected to the feed-back experiments and representing additional 6,770,434 sequencing reads revealed a marked effect of the heat treatment on both the richness and evenness of the rhizosphere communities profiled at the end of cultivation (Kruskal–Wallis followed by a post-hoc Dunn’s test, Individual *P* values < 0.05; Figure S8). Likewise, we observed a compositional shift between heat-treated and native samples (Permanova, R_Treatment_^2^ = 0.289, *P* value = 0.0002, 5,000 permutations, Figure 7C). At the end of cultivation, Bulk soil communities could be separated along the axis accounting for the second source of variation, while the taxonomic composition of rhizosphere samples appeared “to converge” to common profiles. Congruently, when we inspected for the individual bacteria underpinning this diversification, we identified a phylogenetically diverse group of 10 ASVs, whose enrichment in native samples, accounting for ∼10% of the sequencing reads, discriminated between these latter and both unplanted soil and heat-treated samples (Wald test, *P* value <0.05, FDR corrected, Figure 7D).

Taken together, our data suggest that the heat treatment of the soil substrate led to a scenario comparable to a dysbiosis (43) of the rhizosphere microbiota, defined by a low-diversity community composition associated to a reduced growth of the plant host. As the median aboveground dry weight recorded in the feedback experiment is comparable to the one recorded for barley plants grown for the first time in Quarryfield soil (compare data of Figure 7B with the ones of Figure 1), the heat treatment appears to disrupt the capacity of Elite varieties to assemble a taxonomically diverse bacterial consortium associated to optimum barley growth.

## Discussion

Our investigation revealed that nitrogen-availability for plant uptake impacts on the magnitude of the host ‘genotype effect’ on the barley rhizosphere, measured as number of ASVs differentially recruited between genotypes. This effect was maximal at N0%, when measurements of residual N in the rhizosphere were 0 mg Kg^-1^, while it was nearly “obliterated” at N100%, with ∼200 mg Kg^-1^ of residual NO_3_ in the rhizosphere. This is reminiscent of the observation that in *Medicago truncatula*, a model legume, the host control on the microbiota appear exerted in a nitrogen- and genotype-dependent manner (44). Although the modulation of the rhizosphere microbiota in legumes has been associated to plant genes implicated in the establishment of symbiosis with N_2_-fixing bacteria rather than nitrogen nutritional status *per se* (45, 46), it is conceivable that this latter impacts on, at least in part, host-microbe interactions in the barley rhizosphere. This would be congruent with observation gathered from in rice, a cereal phylogenetically related to barley, in which the nitrate transporter *NRT1.1B*, emerged as a critical regulator of both nitrogen use efficiency and microbiota recruitment (47). Likewise, a regulator of lateral root development, designated *LRT1*, emerged as a determinant of microbiota recruitment in maize plants exposed to limiting nitrogen supplies (48).

Rhizodeposition, i.e., the plant-mediated release of organic compounds in the rhizosphere, may represent the nexus between plant’s adaptation to limiting nitrogen supply and microbiota recruitment (16). Consistently, the availability of organic carbon in barley rhizodeposits is inversely correlated with the amount of nitrate concentration in shoots (49). As wild and domesticated barley plants display differential responses to nitrogen fertilisation (50) and a genotype-dependent control of rhizodeposition (51), the characterisation of primary and secondary metabolites released in the barley rhizosphere may provide mechanistic insights into microbiota diversification in barley. However, as recent investigations revealed that the host genetic control of the rhizosphere microbiota in wild and domesticated barley display a quantitative inheritance (32, 34), additional experiments with dedicated genetic material are required to untangle the molecular mechanisms linking nitrogen availability with microbiota diversification.

The observation that the magnitude of the host control on the microbiota was greater when plants were exposed to a nitrogen supply limiting barley growth motivated us to embark on metagenomic survey of this condition. This approach revealed that the microbial communities proliferating at the barely root-soil interface are largely dominated by bacteria: more than 98% of the annotated sequences at phylum level were classified as bacteria. This is strikingly similar to a previous investigation conducted in a soil with different physical and chemical characteristics (30). The dominance of bacterial sequences over other members of the microbiota is not unusual in soil metagenomes (52), although both the protocol used for microbial DNA preparation and the databases used for sequencing annotation (53) can ‘artificially inflate’ the proportion of bacteria among the analysed metagenomes. Despite this potential caveat, we demonstrated with an independent quantification that bacterial gene copy number exceeded that from fungal sources by several orders of magnitude, as previously reported (54, 55). It is however important to consider that PCR-based approaches may fail to provide an accurate estimation of fungal biomass due to nuclear exchanges on these filamentous microorganisms (56).

A closer inspection of the bacterial and fungal abundances revealed no significant differences among samples for Arbuscular Mycorrhizal Fungi (AMF) symbionts of barley. This apparent discrepancy, in view that AMF can mobilise nitrogen for plant uptake (57), is however congruent with a previous investigation conducted with field-grown barley plants, where no genotype effect on AMF root colonisation was observed regardless of the nitrogen regime (58). Furthermore, this observation is also congruent with the fact that our N0% treatment was associated with a replete amount of phosphorus for barley growth, a condition known to suppress AMF colonisation (59). Finally, it is important to mention that the microbiota inhabiting soils whose pH is below 7, such as ‘Quarryfield’, are less conducive to AMF activity than those inhabiting neutral to alkaline substrates (60).

For these reasons, we decided to focus on the functional characterisation of the bacterial component of the microbiota of plants exposed limiting nitrogen supplies. This allowed us to identify three main GO categories enriched in and differentiating rhizosphere samples from bulk soil, namely, ‘carbohydrate metabolic process’, ‘response to chemical’ and ‘pathogenesis’. Of interest is the GO category ‘carbohydrate metabolic process’, whose enrichment emerged as both microhabitat- and genotype-dependent, this is congruent with previous observations that root-derived dissolved organic carbon and carbohydrate utilisation by soil microbes display a host genetic component in wild and domesticated barely genotypes (32, 51). Likewise, ‘response to chemical’ may mirror the adaptation of rhizosphere communities to plant secondary metabolites, released through rhizodeposition, capable of selectively impacting on microbial proliferation, as observed in barley (61) as well as in other cereals (62, 63).

Conversely, the GO category ‘pathogenesis’ appears difficult to reconcile with the fact that no obvious symptoms of disease were observed in our samples. However, studies conducted with the model plant *A. thaliana* revealed that components of host immune system are required for the establishment of a diverse and functional microbiota at the root-soil interface (64) : this suggests that endogenous barley microbiota has evolved the capacity of modulating host immune responses to colonise the rhizosphere. This scenario appears further corroborated by the enrichment of the GO category ‘bacterial-type flagellum-dependent cell motility: despite the fact that molecular components of this machinery have been considered a paradigmatic epitope of the plant immune system (65), it is now emerging that their recognition by host plants contribute to signal modulation and microbiota establishment (66). Similarly, the enrichment of the GO category ‘response to reactive oxygen species’ in the microbiota of the Elite plants may further corroborate the role of this class of compounds in modulating plant-associated bacterial communities (67). A prediction of these observations is that components of the barley immune system may act as a ’checkpoint’ for the taxonomic and functional composition of the rhizosphere microbiota.

Substrate availability and inter-organismal relationships appear to be a determinant also for the bulk soil communities, as mirrored by the significant enrichment of the GO terms ‘photosynthesis’, ‘antibiotic biosynthetic process’ and ‘sporulation’. The absence of a source of organic compounds such as rhizodeposition creates a niche for the proliferation of CO_2_-fixing microorganisms, which are ubiquitous in the soil ecosystem (68). Likewise, the enrichment of ‘antibiotic biosynthetic process’ is congruent with what was observed for agricultural soils in a cross-microbiome survey (69) while sporulation underpins microbial adaptation to soil stressful conditions (70). Furthermore, unplanted soil communities display a differential enrichment for function implicated in phosphorous homeostasis. As the relative abundances of carbon, nitrogen and phosphorus can be considered constrained in microbial biomass (71), this observation suggests that, although phosphorous was applied with the nutrient solution to all specimens, this element may act as limiting factor predominantly for unplanted soil communities, where the lack of exudates reduce phosphorous solubility.

Taken together, these observations provide mechanistic insights into the multi-step selection process differentiating rhizosphere communities from bulk soil ones (4, 27) implicating the modulation of host immune responses as one of the requirements for bacterial establishment in the rhizosphere of plants exposed to limiting nitrogen supply. However, as these experiments were performed in a single soil type, caution is required in extrapolating the results as being indicative of general phenomena applicable across all soils. Further metagenomics investigations with plants exposed to replete nitrogen conditions, benefiting also from latest development in sequencing technologies (72), be required to accurately gauge the impact of this mineral (or lack thereof) on the functional potential of the barley microbiota.

Despite the fact that the 60 MAGs generated in this work accounted for less than 10% of the metagenomic reads, these figures are aligned with what has been recently observed for the rhizosphere of sorghum (73), a cereal phylogenetically related to barley. This effort not only allowed us to identify genomes belonging to the dominant phyla of the plant microbiota (i.e., Actinobacteria, Bacteroidetes, Firmicutes and Proteobacteria) but also members of additional classes, such as an individual member of the metabolically diverse and yet poorly characterised Zixibacteria phylum (74-76). Furthermore, mapping reads associated to the GO terms differentially enriched between microhabitat and genotype, allowed us to gain novel insights into the relationships between taxonomic and functional composition of the barley microbiota. For instance, we observed an association between the GO category ‘Carbohydrate metabolic process’ and the enrichment of members of the phylum Bacteroidetes in the ‘Elite’ rhizosphere. As cell wall features represent a recruitment cue for the plant microbiota (77), this enrichment may mirror the capacity of degrading complex polysaccharides coded by members of this phylum (78, 79). The observed genotype-specific enrichment may be further explained by polymorphisms of barley genes regulating carbohydrate composition in the cell wall (80, 81).

The plant-soil feedback experiment we implemented suggested that, a functional rhizosphere microbiota is required for optimal barley growth under nutrient limiting conditions. Although not significantly different, mean values of aboveground biomass of Elite plants recorded in the ‘Desert’-conditioned soil were higher than recorded from the soil conditioned with the same genotype. Despite phylogenetic relatedness between condition and focal species in plant-soil feedback experiments appear unrelated to the strength of the feedback itself (82, 83), compositional shifts between the conditioned and focal microbiota tend to be associated with enhanced plant growth (84). However, significant differences in growth were observed when Elite plants were exposed to heat-inactivated soils which are associated to a reduction of alpha diversity indices in the rhizosphere, a condition which has previously linked to stressful soil conditions (84). In turn, this effect could be due to the treatment on the microbes per se, release of mineral nutrients and/or the disruption liable carbon compounds released through exudates (85) by conditioning plants and capable of modulating individual members of the barley microbiota (86). Unlike recent observations gathered from plant-soil feedback experiments of maize plants exposed to limiting nitrogen conditions (48), what emerged from our study is the control exerted by the recipient genotype on the resulting bacterial communities. This was manifested by the microbiota of plants exposed to either Desert-conditioned or Elite-conditioned soil converging towards a phylogenetically conserved bacterial consortium. This is in accordance with data gathered from rice, using both soil feedback experiments (87) and synthetic communities (47), indicating the host genotype as a driver of a plant-growth promoting microbiota. Likewise, a recently developed indexed bacterial collection of the barley rhizosphere microbiota indicated a growth-promotion potential for members of the phyla differentially recruited in the feedback experiment (88, 89).

Taken together, this suggests that the enriched bacteria represent a consortium of beneficial bacteria required for optimum barley growth whose recruitment is driven, at least in part, by the host genotype.

### Conclusions

Our results point at nitrogen availability for plant uptake as inversely correlated with the magnitude of the host genetic control on the taxonomic composition of the barley rhizosphere microbiota. Under nitrogen supply limiting barley growth, wild and domesticated genotypes retain specific functional signatures which appear to be encoded by distinct bacterial members of the microbiota. Although we found evidence for nitrogen metabolism executed by these communities, adaptation to the plant immune system emerged as an additional recruitment cue for the barley microbiota. Plant-soil feedback experiments suggest that these distinct compositional and functional configurations of the microbiota can be “rewired” by the host genotype leading to a recruitment of a consortium of bacteria putatively required for optimum plant growth. Thanks to recent insights into barley genes shaping the rhizosphere microbiota (32, 34), these concepts can now be tested under laboratory and field conditions to expedite the development of plant varieties combining profiting from improved yields with reduced impacts of N-fertilisation on the environment.

## Methods

### Experimental design

This is investigation consists of three distinct but interconnected experiments. For each one of them, plants were maintained under controlled conditions in the same soil type designated ‘Quarryfield’ (see ‘Soil’ below) and individual samples were arranged in a completely randomised design. For the first experiment, we grew and subjected to 16S rRNA gene amplicon sequencing individual biological replicates (i.e., pots) Elite, Desert and North as well as three Bulk soil controls exposed to three different nitrogen treatments, designated N0%; N25% and N100% respectively (see ‘Nitrogen treatments’ below) according to the following scheme. N0%, number of sequenced replicates for Desert N0%_Desert_ = 5; N0%_North_ = 3; N0%_Elite_ = 4; N0%_Bulk_ = 4. N25%, number of sequenced replicates for Desert N25%_Desert_ = 5; N25%_North_ = 3; N25%_Elite_ = 4; N25%_Bulk_ = 4. N100%, number of sequenced replicates for Desert N100%_Desert_ = 5; N100%_North_ = 4; N100%_Elite_ = 3; N100%_Bulk_ = 4. Alongside these samples, we prepared two additional Bulk soil controls amended with a plug of the agar substrate used for seed germination. Total number of sequenced samples = 50. In the second experiment, we grew and subjected to shotgun metagenomic sequencing three individual biological replicates (i.e., individual plants in individual pots) of the genotypes Elite, Desert and North as well as three Bulk soil controls exposed to N0% treatment. Total number of sequenced samples = 12. In the third and final experiment, we grew and subjected to 16S rRNA gene amplicon sequencing individual biological replicates (i.e., individual plants in individual pots) of the Elite genotype soil controls in Quarryfield soils previously conditioned (see ‘Plant-soil feedback experiment’ below) with either the Elite or Desert genotype in a native form or upon heat treatment. For the former, we contemplated also Bulk soil control pots. Upon discarding pots with no detectable plant growth, the number of rhizosphere samples exposed to Elite-conditioned soil retained for sequencing was Elite-native_rhizosphere_ = 11; Elite-native_Bulk_= 7. Desert-conditioned soil, Desert-native_rhizosphere_ = 14; Desert-native_Bulk_= 9. Number of sequenced rhizosphere samples exposed to the heat treated soil, Elite-treated_rhizosphere_ = 15; Desert-treated_rhizosphere_= 15. Total number of sequenced samples = 71.

### Soil

Soil was sampled from the agricultural research fields of the James Hutton Institute, Invergowrie, Scotland, UK in the Quarryfield site (56°27’5“N 3°4’29”W). This is a sandy silt loam soil with a pH of 6.2 and 5% organic matter content. The nitrogen content of this soil was 1.8 mg Kg^-1^ ammonium and 13.5 mg Kg^-1^ nitrate. The site was left unplanted and unfertilised in the three years preceding the investigations.

### Plant material and growth conditions

Barley seeds of the domesticated (*Hordeum vulgare* ssp. *vulgare*) and wild (*Hordeum vulgare* ssp. *spontaneum*) genotypes, the variety ‘Morex’ (i.e., ‘Elite’) and the accessions B1K-12 (i.e., ‘Desert’) and B1K-31 (i.e., ‘North’), respectively, were surface sterilized as previously reported (90) and germinated on 0.5% agar plates at room temperature. Seedlings displaying comparable rootlet development were sown individually in 12-cm diameter pots containing approximately 500g of the ‘Quarryfield’ soil, previously sieved to remove stones and large debris. Unplanted pots filled with the same soil, i.e., bulk soil controls, were maintained in the same glasshouse and subjected to the same treatments as planted pots. Plantlets one week old were transferred for two weeks to a growth room at 4 °C for vernalisation. Following the vernalisation period, plants were maintained in a randomized design in a climatic-controlled glasshouse at 18/14 °C (day/night) temperature regime with 16 h daylight that was supplemented with artificial lighting to maintain a minimum light intensity of 200 µmol quanta m^−2^ s^−1^ until early stem elongation (Supplementary Figure 1). Watering was performed weekly as indicated (see ‘Nitrogen treatments’ below). Pots were rotated on weekly basis to minimise potential biases associated to given positions in the glasshouse.

### Nitrogen treatments

The nutrient solutions described in this study, i.e., N100%, N25% and N0% are reported in Supplementary Table 1. Nutrient solutions were applied at a rate of 25ml per kg of soil each week. Applications started two days after planting, were interrupted during the vernalisation and reinstated once the plants were transferred to the growing glasshouse and they reached early stem elongation. Fourteen treatments were applied with a total of 312.5 mg (NO_3_ ^-^); 81.4 mg (NH_4_ ^+^) for the N100% solution, 78.1 mg (NO_3_ ^-^); 20.0 mg (NH_4_ ^+^) for the N25% solution, and 0 mg of (NO_3_ ^-^ and NH_4_ ^+^) for the N0% solution per pot.

### Plant and Soil Nitrogen determination

To assess the N content of the plant, at the time of sampling a newly expended leaf was sectioned from every plant, freeze-dried, ball milled, and N content measured in an Elemental Analyser CE-440 (Exeter Analytical Inc, UK). The soil from the pots was sieved through a 2mm mesh sieve and mixed. Five grams of soil was added to 25mL of 1M KCl and the resulting solution mixed in a tube roller for 1hour at ∼150 rpm. Supernatant was transferred to 50mL falcon tubes and centrifuged for 15min at 5,000 rpm, then the supernatant was subject to another round of centrifugation. The supernatant was transferred to a falcon tube and analysed with a Discrete Analyser Konelab Aqua 20 (Thermo Fisher, Waltham, USA) in the analytical services of The James Hutton Institute (Aberdeen, UK). In parallel, ∼10 g from the sieved soil was oven dried at 70 °C for 48h and dry weight recorded to express the analytical results in NO_3_ ^-^ and NH_4_ ^+^ in mg N kg^-1^ of soil.

### Bulk soil and rhizosphere DNA preparation

At early stem elongation, plants were excavated from the soil and the stems were separated from the roots. The uppermost 6 cm of the root system were detached from the rest of the root corpus and processed for further analysis. The sampled aboveground material was oven dried at 70°C for 48 hours and the dry weight recorded. The roots were shaken manually to remove loosely attached soil. For each barley plant, the seminal root system and the attached soil layer was collected and placed in a sterile 50ml falcon tube containing 15ml phosphate-buffered saline solution (PBS). Rhizosphere was operationally defined, for these experiments, as the soil attached to this part of the roots and extracted through this procedure. The samples were then vortexed for 30 seconds and transferred to a second 50ml falcon containing 15ml PBS and vortexed again to ensure the dislodging and suspension of the rhizosphere. Then, the two falcon tubes with the rhizosphere suspension were combined and centrifuged at 1,500g for 20 minutes to precipitate the rhizosphere soil into a pellet, then flash frozen with liquid nitrogen and stored at -80°C until further analysis. In addition, we incubated water agar plugs (∼1 cm^3^) into two unplanted soil pots and we maintained them as control samples among the experimental pots to monitor the effect of this medium on the soil microbial communities. DNA was extracted from unplanted soil and rhizosphere samples using FastDNA™ SPIN Kit for Soil (MP Biomedicals, Solon, USA) according to the manufacturer’s recommendations and stored at -20°C.

### Preparation of 16 rRNA gene amplicon pools

The hypervariable V4 region of the small subunit rRNA gene was the target of amplification using the PCR primer pair 515F (5’-GTGCCAGCMGCCGCGGTAA-3’) and 806R (5’-GGACTACHVGGGTWTCTAAT-3’). The PCR primers had incorporated an Illumina flow cell adapter at their 5’ termini and the reverse primers contained 12bp unique ‘barcode’ for simultaneous sequencing of several samples (91). PCR reactions were performed using 50 ng of metagenomic DNA per sample using the Kapa HiFi HotStart PCR kit (Kapa Biosystems, Wilmington, USA). The individual PCR reactions were performed in 20 µL final volume and containing 4 µL of 5X Kapa HiFi Buffer, 10 µg Bovine Serum Albumin (BSA) (Roche, Mannheim, Germany), 0.6 µL of a 10 mM Kapa dNTPs solution 0.6 µL of 10 µM solutions of the 515F and 806R PCR primers and 0.25 µL of Kapa HiFi polymerase. The reactions were performed using the following programme programme: 94 °C (3 min), followed by 35 cycles of 98 °C (30 s), 50 °C (30 s) 72 °C (1 min) and a final step of 72 °C (10 min). For each primer combination, a no template control (NTC) was included in the reactions. To minimise amplification biases, PCRs were performed in triplicate and at least two independent master mixes per barcode were generated (i.e., 6 reactions/sample). PCR reactions were pooled in a barcode-wise manner and an aliquot of each amplification product inspected on 1.5% agarose gel. Only samples whose NTCs yielded an undetectable PCR amplification were retained for further analysis. PCR purification was performed using Agencourt AMPure XP Kit (Beckman Coulter, Brea, USA) with 0.7 µL AmPure XP beads per 1 µL of sample. Following purification, each sample was quantified using PicoGreen (Thermo Fisher Scientific, Watham, USA) and individual barcode samples were pooled in an equimolar ratio to generate amplicons libraries.

### Illumina 16S rRNA gene amplicon sequencing

The pooled amplicon library was submitted to the Genome Technology group, The James Hutton Institute (Invergowrie, UK) for quality control, processing and sequencing. Amplicon libraries were amended with 15% of a 4pM phiX solution. The resulting high-quality libraries were run at 10 pM final concentration on an Illumina MiSeq system with paired-end 2x 150 bp reads (91) to generate the sequencing output, the FASTQ files.

### Amplicon sequencing reads processing

Sequence reads were subjected to quality assessment using FastQC (92). ASVs were then generated using DADA2 version 1.10 (93) and R 3.5.1 (94) following the basic methodology outlined in the ‘DADA2 Pipeline Tutorial’ (95).Read filtering was carried out using the DADA2 paired FastqFilter method, trimming 10bp of sequence from the 5’ of each read using a truncQ parameter of 2 and maxEE of 2. The remainder of the reads were of high quality consequently no 3’ trimming was deemed necessary. The dada2::learn_errors() method was run to determine the error model with a MAX_CONSIST parameter of 20, following which the error model converged after 9 and 12 rounds for the forward and reverse reads respectively. The dada2::dada() method was then run with the resulting error model to denoise the reads using sample pooling, followed by read merging, followed by chimera removal using the consensus method. Taxonomy assignment was carried out using the RDP Naive Bayesian Classifier through the dada2::assignTaxonomy() method, with the SILVA database (96) version 138, using a minimum bootstrap confidence of 50. The DADA2 outputs were finally converted to a Phyloseq object (version 1.26.1) (97).

The Phyloseq objects for both the nitrogen gradient and the Plant-soil feedback experiments were initially merged. Next sequences classified as either ‘Chloroplast’ or ‘Mitochondria’ were pruned in silico from the merged object. Likewise, ASVs matching a list of potential contaminants of the lab (98) were removed as well as ASVs lacking a taxonomic classification at phylum level (i.e., ‘NA’). We further applied an abundance filtering and retained ASVs occurring with at least 20 reads in 2% of the samples. Finally, the Phyloseq objects were rarefied at 25,000 reads per sample, as recommended for groups with large differences in library size (99), prior to downstream analyses.

### Metagenome sequencing, annotation and analysis

We generated a new set of 3 bulk soil, and 3 rhizosphere DNA preparations from each of the three genotypes tested (i.e., ‘Desert’, ‘North’ and ‘Elite’) from specimens maintained in Quarryfield soil under N0% conditions as described above. These 12 new preparations were quantified and submitted to the LGC Genomics sequencing service (Berlin, Germany) where they were used to generate DNA shotgun libraries using the Ovation Rapid DR Multiplex System 1-96 NuGEN (Leek, The Netherlands) kit following manufacturer’s recommendations. These libraries were run simultaneously into an individual Illumina NextSeq500 run following manufacturer’s recommendations with the 2 X150bp chemistry and generated a total of 412,385,413 read pairs. After sequencing read pairs were de-multiplexed according to the sample’s barcodes using the Illumina bcl2fastq2.17.1.14 software.

Metagenome analysis was conducted according to the general approach of Hoyles *et al* (100), using updated tools where appropriate. Sequence reads were quality assessed using FastQC (101) and quality/adapter trimmed using TrimGalore (102), using a quality cut-off of 20, a minimum sequence length of 75bp and removing terminal N bases. Taxonomic classification of the sequence reads was carried out using Kraken 2.0.9 (103) with the Kraken PlusPFP database (104), which incorporates protozoa, fungi and plants in addition to the archaea, bacteria and viruses present in the standard database. Host contamination was removed by alignment against the Morex V2 barley genome sequence (105) using BWA MEM (106), and non-aligning reads extracted from the resulting bam files using SAMtools

(107). Metagenome assembly was conducted using MegaHit version 1.2.9 (108) with the ‘meta-large’ preset. Predicted proteins were produced from all assemblies using Prodigal version 2.6.3 (109), which were then clustered using MMseqs2 version 11.e1a1c (110) using the ‘easy_cluster’ method. Abundance of predicted proteins in each sample was determined by alignment of sequence reads against the representative cDNA sequences of the clusters using BWA MEM and determining the read counts associated with each sequence using a custom PySAM (111) script. Functional annotations of the protein sequences were carried out using InterProScan 5-50-84.0 (112) and Interpro version 84.0. GO terms were enumerated using a custom python script which assessed the number of occurrences of each term in each sample based upon the previously determined abundance of each annotated sequence. GO terms were mapped to the metagenomics GO slim subset dated 2020-03-23 (113) using the Map2Slim function of OWLtools (114). Functional enrichment analysis was carried out using DESeq2 (115) version 1.26.0.

### Metagenome-Assembled Genomes (MAGs)

MAGs were created using the MegaHit-assembled contigs described above using MetaBat2 version 2.15 to create contig bins representing single genomes. Contig bins were dereplicated using dRep version 3.2.0 followed by decontamination with Magpurify version 2.1.2 (116). The resulting MAGs were assessed for completeness and contamination using checkM (117). Annotation of the MAGs was carried out with Prokka 1.14.6 (118) and InterProScan 5-50-84.0 (112), before taxonomic classification was determined using GTDB-TK version 1.4.0 (119) with data version r95.

### Nitrogen Cycle Gene Analysis

Abundance of nitrogen-cycle genes was determined using the NCycProfiler tool of NCycDB (120) with the diamond method. Pairwise t-tests were carried out between the samples of each group within each gene to identify combinations with statistical differences between samples (Benjamini-Hochberg corrected, FDR<0.05).

### Plant-soil feedback experiment

We grew Desert and the Elite genotypes in ‘Quarryfield’ soil supplemented with a N0% nutrient solution under controlled conditions (see Plant growth conditions). We selected these two genotypes since, in the tested soils, they host a taxonomic and functional distinct microbiota. At early stem elongation we removed the plants from the soil, and we harvested the residual soil and kept it separated in a genotype-wise manner. We reasoned that at the end of cultivation the soils would have been enriched, at least partially, for specific microbial taxa and functions associated to either genotype. This residual soil, either in a ‘native form’, i.e., not further treated after sampling, or after being exposed to a heat-treatment (126°C for 1 hour, repeated twice at an interval of ∼12 hours), was used as a substrate for a subsequent cultivation of a recipient Elite barley genotype. These plants were maintained under controlled conditions (see Plant growth conditions) and supplemented with a N25% solution to compensate for the near complete depletion of this mineral in the previous cycle of cultivation (compare the NH_4_ ^+^ and NO_3_ ^-^ concentrations of rhizosphere specimens at N0% and N25% in Figure 1). At early stem elongation plants were harvested and their aboveground biomass determined after drying stems and leaves at 70°C for 48 hours. At the end of each replicated experiment, the residual soil was collected and subjected to chemical and physical characterisation (Yara United Kingdom Ltd., Grimsby, United Kingdom).

A quantitative real-time polymerase chain reaction assay was used to quantify the bacterial and fungal DNA fractions in samples from the conditioned soil experiment as follows. DNA samples were diluted to 10 ng/µl and successively diluted in a serial manner to a final concentration of 0.01 ng/µl. This final dilution was used for both the Femto Fungal DNA Quantification Kit and Femto Bacterial DNA Quantification Kit (Zymo Research) and the quantification was conducted according to the manufacturers protocol. Briefly, two microliters of the 0.01 ng/µl dilution of each sample was used together with 18 µl of the corresponding fungal or bacterial master mix. Two µl of the fungal or bacterial standards were also used to create the respective quantification curves. DNA samples from the conditioned soil experiment were randomized in the 96 well plates, using a minimum of 11 biological replicates per treatment. The quantification was performed in a StepOne thermocycler (Applied Biosystems by Life Technology) following the cycling protocols of each of the above-mentioned bacterial and fungal kits.

### Statistical analyses on univariate dataset and amplicon sequencing

Data analysis was performed in R software using a custom script with the following packages: Phyloseq (97) version 1.30.0 for pre-processing, alpha and beta-diversity analysis; ggplot2 version 3.3.0 (121) for data visualisations; Vegan version 2.5-6 (122) for statistical analysis of beta-diversity; PMCMR version 4.3 (123) for non-parametric analysis of variance and Agricolae for Tukey post hoc test (124). For any univariate dataset used (e.g., aboveground biomass) the normality of the data’s distribution was checked using Shapiro–Wilk test. For datasets normally distributed, the significance of the imposed comparisons was assessed by an ANOVA test followed by a Tukey post hoc test. Non-parametric analysis of variance were performed by Kruskal-Wallis Rank Sum Test, followed by Dunn’s post hoc test with the functions kruskal.test and the posthoc.kruskal.dunn.test, respectively, from the package PMCMR. We used Spearman’s rank correlation to determine the similarity between unplanted soil profiles and bulk soil samples amended with water agar plugs (Table S2). The analysis of ASVs differentially enriched was performed a) between individual genotypes and bulk soil samples to assess the sample effect and b) between the rhizosphere samples to assess the genotype effect. The genotype effect was further corrected for a microhabitat effect (i.e., for each genotype, only ASVs enriched against both unplanted soil and at least another barley genotype were retained for further analysis). The analysis was performed using the DESeq2 package (115) version 1.26.0 consisting of a moderated shrinkage estimation for dispersions and fold changes as an input for a pair-wise Wald test. This method identifies the number of ASVs significantly enriched in pair-wise comparisons with an adjusted *P* value (False Discovery Rate, FDR < 0.05). This method was selected since it outperforms other hypothesis-testing approaches when data are not normally distributed and a limited number of individual replicates per condition are available (99).

## Supplemental Material

**Table S1**. Composition of the nutrient solutions used in this study. The solution was applied with watering of the plants at a rate of 25 ml of the nutrient solution per Kg of soil.

**Table S2**. Spearman’s rank correlations computed between the average relative abundances (phylum level) of the communities retrieved from unplanted soil samples and unplanted soil amended with “0.5% agar plugs”.

**Figure S1: Barley development at the time of sampling.** Representative photographs of the indicated genotypes subjected to different nitrogen treatment taken at the time of sampling. Scale bar = 5 cm.

**Figure S2: Schematic representation of the metagenomics computational pipeline. Figure S3: Nitrogen biogeochemical cycle genes at the barley root-soil interface**. Individual panels depict metagenomics sequencing reads assigned to a given gene. Dots depict individual biological replicated color-coded according to sample affiliation as indicated at the bottom of the panels. Only genes displaying a significant difference between samples are presented (pairwise t-tests, individual P value <0.05, BH corrected). Individual gene abbreviations: gdh, Glutamate dehydrogenase; hao, Hydroxylamine dehydrogenase; napA, Periplasmic nitrate reductase; narB Assimilatory nitrate reductase; narC Cytochrome b-561; nirA, Ferredoxin-nitrite reductase; nirB, Nitrite reductase (NADH) large subunit; norB, Nitric oxide reductase subunit B; nosZ, Nitrous-oxide reductase; NR, Nitrate reductase (NAD(P)H); nrfA, Nitrite reductase (cytochrome c-552); ureA, Urease subunit gamma; ureB Urease subunit beta.

**Figure S4: K-Means clustering of GO annotations in metagenomic samples**. These panels indicate the number of rlog-transformed counts for each cluster centroid resulting from K-means clustering (10 clusters, maximum of 40 iterations) which reveals consistent patterns amongst replicates providing a finer-grained view of functional enrichment. Cluster 5, for example contains GO terms which are increased in abundance in both desert and north sample relative to bulk soil, with even higher abundance in elite samples. Members of cluster 6 are similarly increased in abundance in all planted samples relative to bulk soil. Cluster membership, including DESeq2 differential abundance analysis results are presented in Dataset S1.

**Figure S5: MAGs differential enrichment across microhabitats and genotypes.** Each panel denotes a pair-wise comparison between bulk soil and rhizosphere (top panels) or between genotypes within the rhizosphere microhabitat (bottom panels). Differential enrichments expressed a log2 fold change, with enrichment in the first term of comparison depicted by negative fold change (Wald test, Individual P values <0.05, FDR corrected).

**Figure S6: Impact of the heat treatment on soil chemical and physical parameters.** Non-metric multidimensional scaling illustrating the relationships of soil samples from the indicated sample type. Arrows depict the most significant parameters (R^2^ > 0.83, *P* value <0.001) explaining the ordination, namely the concentration of ammonium, phosphorus, manganese, sulphur, sodium, iron, copper, zinc and the pH. Arrows point at the direction of change while their length is proportional to the correlation between the ordination and the indicated variables.

**Figure S7: Bacterial and fungal DNA concentration in samples from the plant-soil feedback experiment.** Boxplots depicting the logarithm (base 2) of the concentration (expressed as copy numbers per 2μl of input DNA) of the A) 16 rRNA gene or B) ITS sequences retrieved from the plant-soil feedback experiment. Individual dots depict individual biological replicated. Different letters denote significantly different groups (Kruskal–Wallis and post-hoc Dunn’s test, *P* value = 0.039); ns no significant differences.

**Figure S8: The heat treatment impacts on microbiota richness and evenness.** Strip chart depicting A) number of ASVs or B) Shannon indexes of the samples subjected to the plant-soil feedback experiments. Individual dots depict individual biological replicates whose colour reflects the sample type and the treatment indicated at the bottom of the figure. Different letters denote significantly different groups (Kruskal–Wallis and post-hoc Dunn’s test, *P* value < 0.05).

**Dataset S1.** Analysis of individual GO-terms identified on the top 10 clusters differentiating between samples and arranged as individual spreadsheets. For each cluster we determined the significance of individual terms in pair-wise comparisons between bulk soil and rhizosphere samples and, within the latter, between genotypes (Wald test, Individual P values <0.05, FDR corrected).

## Acknowledgements

We are particularly grateful to Dr Eyal Fridman (ARO, Bet Dagan, Israel) for providing us with the B1K seeds used in this study. We thank Lawrie Brown (The James Hutton Institute, Invergowrie) for advising us in the preparation and application of the nutrient solutions, Niamh Johnston (Nuffield Research Placements Student) and Jim Wilde (The James Hutton Institute, Invergowrie) for the technical assistance during the feedback experiment. We thank LGC genomics Gmbh (Berlin, Germany) for generating the shot gun metagenomics reads. For the purpose of open access, the authors haves applied a CC BY public copyright licence to any Author Accepted Manuscript version arising from this submission.

## Availability of data and materials

The sequences generated in the 16S rRNA gene sequencing survey and the raw metagenomics reads reported in this study are deposited in the European Nucleotide Archive (ENA) under the accession numbers PRJEB30847, PRJEB54872 and PRJEB54873. Individual metagenomes are retrievable on the MG-RAST server under the IDs mgm4798244.3; mgm4798274.3; mgm4798349.3; mgm4798388.3; mgm4798507.3; mgm4798563.3; mgm4798641.3; mgm4798894.3; mgm4799467.3; mgm4799972.3; mgm4801514.3; mgm4801719.3.

The scripts used to analyse the data and generate the figures of this study are available at https://github.com/BulgarelliD-Lab/Barley-NT-2020

## Authors’ contribution

The study was conceived by RAT and DB with critical inputs from EP and LB. RAT, AMC, CEM, KBC and RK performed the experiments. JM and PH generated the 16S rRNA gene sequencing reads. JA conceived the metagenomic analysis with inputs from MB and GT. RAT, SRA, AMC, CEM, JA and DB analysed the data. All authors critically reviewed and edited the manuscript and approved its publication.

