## Supplementary figures and images for "Defining composition and function of the rhizosphere microbiota of barley genotypes exposed to growth-limiting nitrogen supplies"

### Supplementary Figure 1

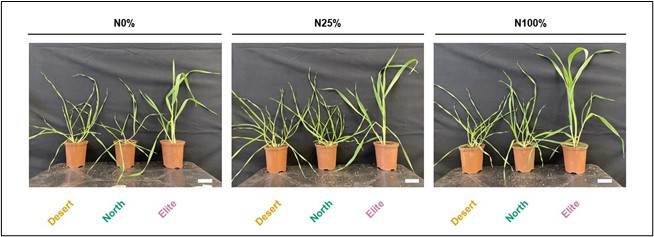

### Supplementary Figure 2

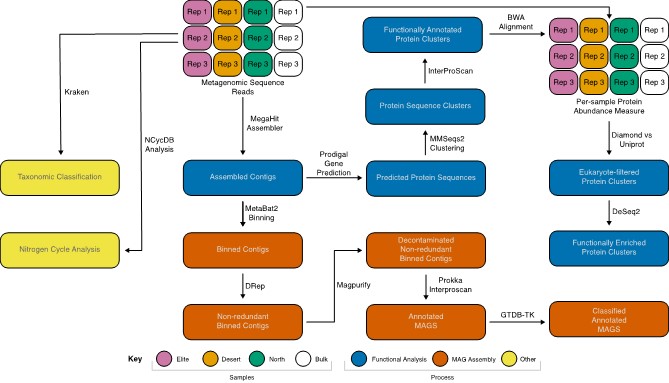

### Supplementary Figure 3

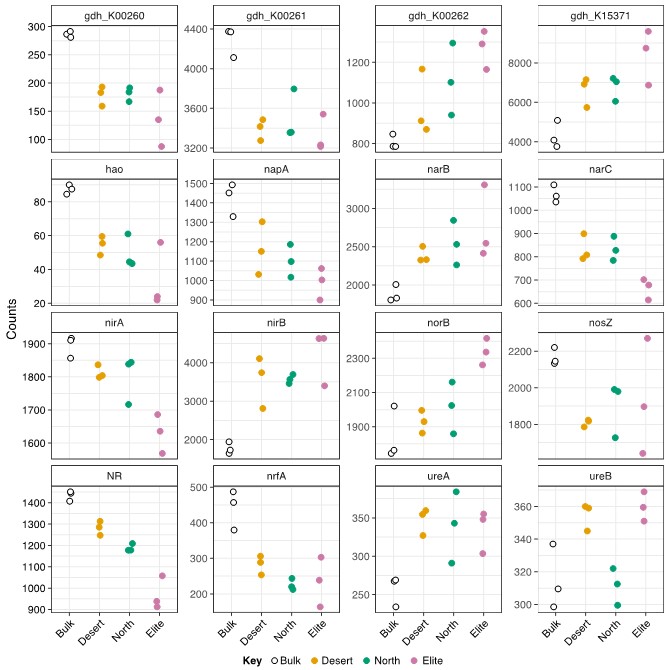

### Supplementary Figure 4

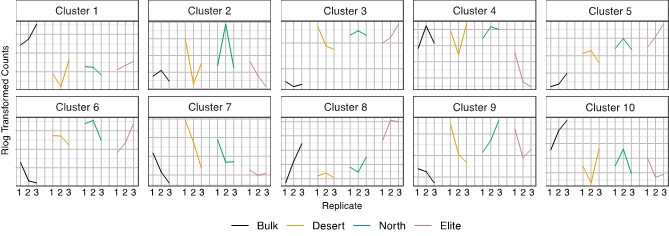

### Supplementary Figure 5

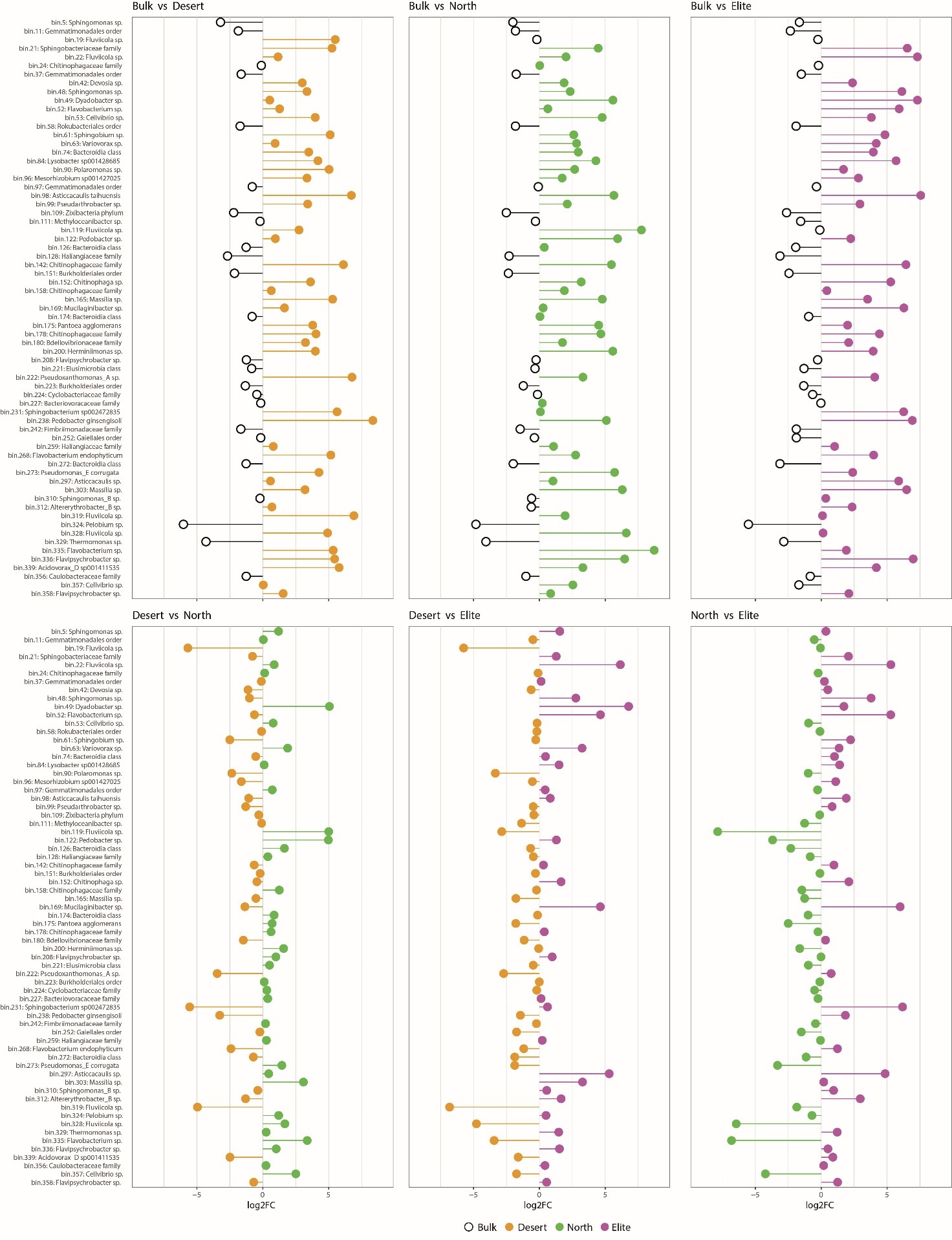

### Supplementary Figure 6

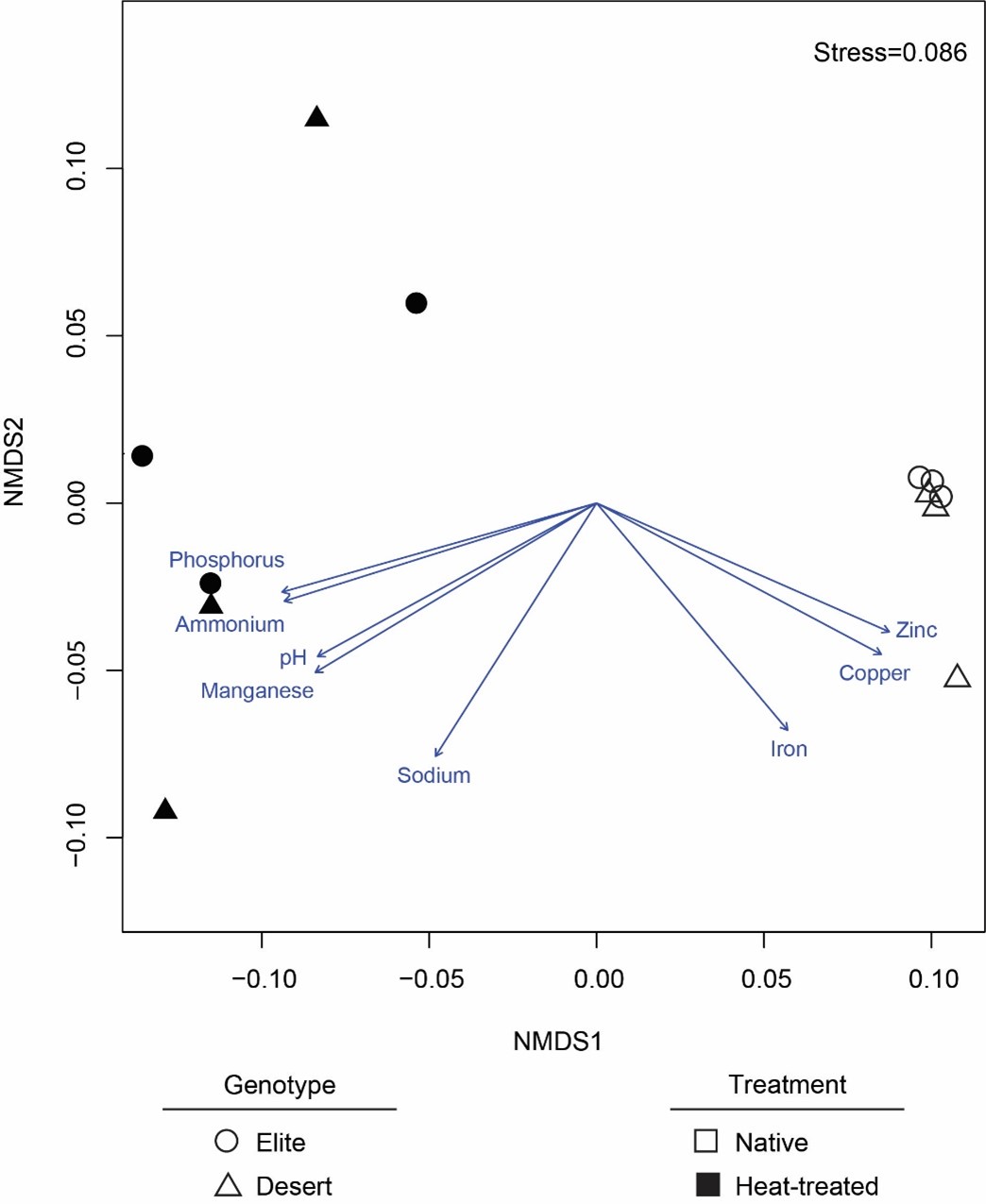

### Supplementary Figure 7

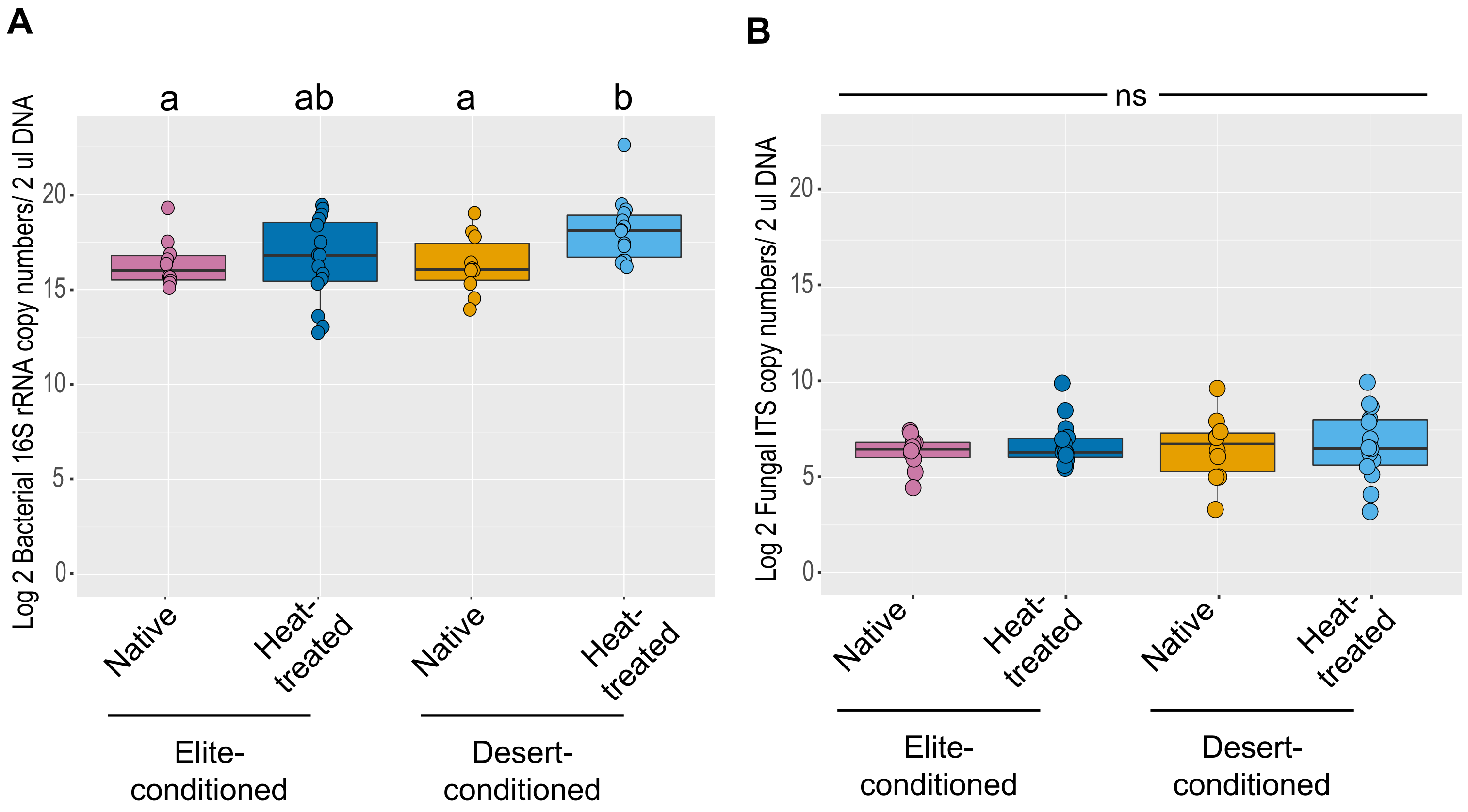

### Supplementary Figure 8

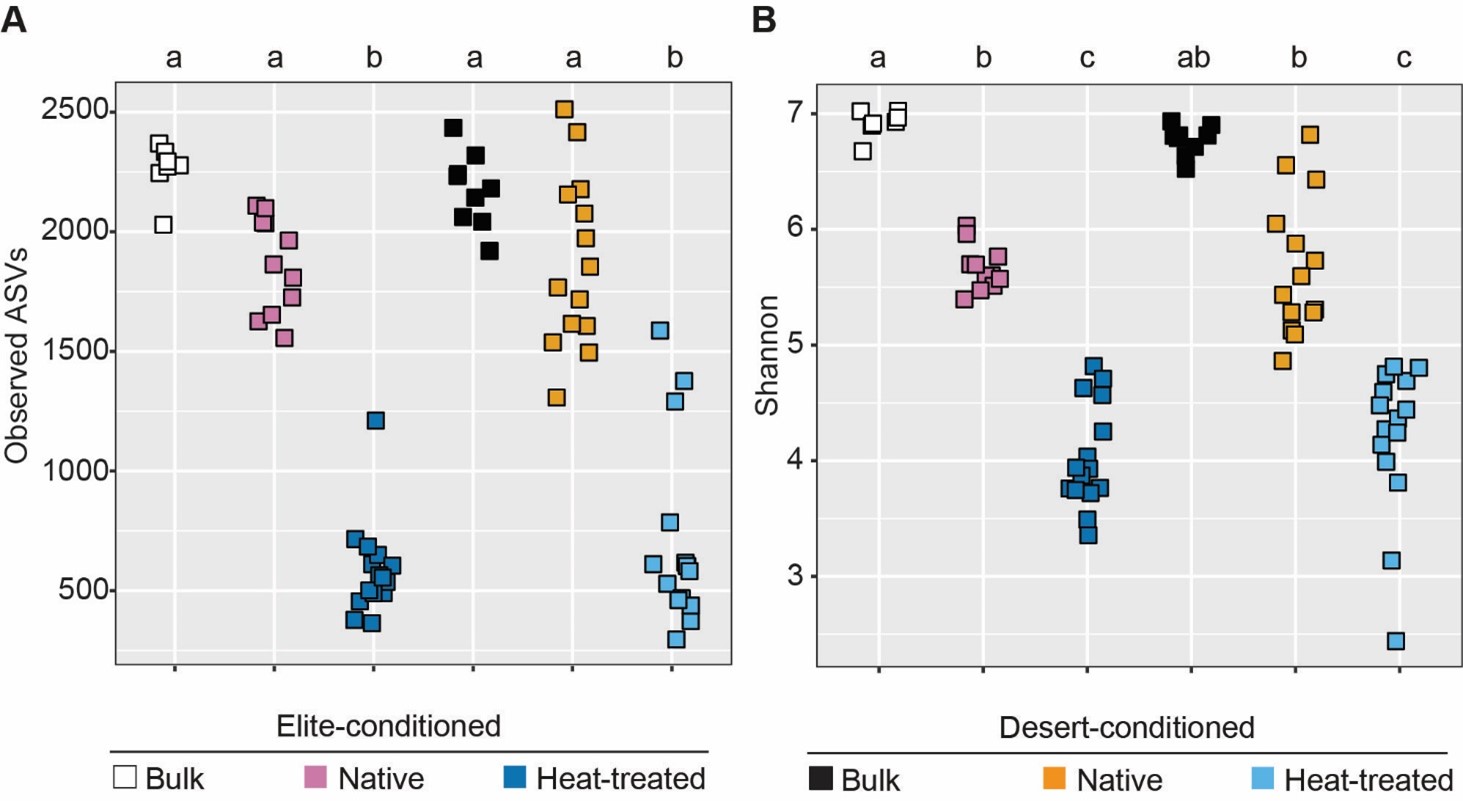
