## Supplementary Table 1 for "Defining composition and function of the rhizosphere microbiota of barley genotypes exposed to growth-limiting nitrogen supplies"

Table S1. Composition of the nutrient solutions used in this study. The solution was applied with watering of the plants at a rate of 25 ml of the nutrient solution per Kg of soil.

|  | **N 100%** | **N 25%** | **N 0%** |
| --- | --- | --- | --- |
| 2(NH_4_) SO_4_ | 25mM | 6.25mM | 0 |
| Ca(NO_3_)_2_ | 40mM | 10mM | 0 |
| KNO_3_ | 10mM | 2.5mM | 0 |
| Mg SO_4_ | 3mM | 3mM | 3mM |
| FeEDTA + | 100μM | 100μM | 100μM |
| Micronutrients* | See below | | |
| KH_2_PO_4_ | 1mM | 1mM | 1mM |
| CaSO_4_ + 2H_2_O | 0 | 0.019mM | 0.025mM |
| CaCl_2_ | 0 | 0.06mM | 0.08mM |
| KCl | 0 | 0.19mM | 0.25mM |

*Micronutrients concentration: 6 μM MnCl_2_, 23 μM H_3_BO_3_, 1.6 μM CuSO_4_, 0.6 μM ZnSO_4_, 1 μM Na_2_MoO_4_, 1 μM CoCl_2_.

|  | Soil water agar controls | |
| --- | --- | --- |
|  | **Spearman’s rho** | **P value** |
| Bulk N0% | 0.9743402 | < 2.2e-16 |
| Bulk N25% | 0.9747067 | <2.2e-16 |
| Bulk N100% | 0.9450147 | < 2.2e-16 |
