## Supplementary Table 2 for "Defining composition and function of the rhizosphere microbiota of barley genotypes exposed to growth-limiting nitrogen supplies"

Table S2. Spearman’s rank correlations computed between the average relative abundances (phylum level) of the communities retrieved from unplanted soil samples and unplanted soil amended with “0.5% agar plugs”.

|  | Bulk amended with 0.5% agar ‘plugs’ | |
| --- | --- | --- |
|  | **Spearman’s rho** | **P value** |
| Bulk N0% | 0.974 | < 2.2e-16 |
| Bulk N25% | 0.975 | <2.2e-16 |
| Bulk N100% | 0.945 | < 2.2e-16 |

|  | Bulk amended with 0.5% agar ‘plugs’ | |
| --- | --- | --- |
|  | **Spearman’s rho** | **P value** |
| Bulk N0% | 0.974 | < 2.2e-16 |
| Bulk N25% | 0.975 | <2.2e-16 |
| Bulk N100% | 0.945 | < 2.2e-16 |
